# VISUALIZING GAUSSIAN-CHAIN LIKE STRUCTURAL MODELS OF HUMAN α-SYNUCLEIN IN MONOMERIC PRE-FIBRILLAR STATE: SOLUTION SAXS DATA AND MODELING ANALYSIS

**DOI:** 10.1101/2024.07.07.602212

**Authors:** Madhumita Dey, Arpit Gupta, Maulik D. Badmalia, Ashish, Deepak Sharma

## Abstract

Here, using small angle X-ray scattering (SAXS) data profile as reference, we attempted to visualize conformational ensemble accessible prefibrillar monomeric state of α-synuclein in solution. In agreement with previous reports, our analysis also confirmed that α-synuclein molecules adopted disordered shape profile under non-associating conditions. Chain-ensemble modeling protocol with dummy residues provided two weighted averaged clusters of semi-extended shapes. Further, Ensemble Optimization Method (EOM) computed mole fractions of semi-extended “twisted” conformations which might co-exist in solution. Since these were only C^α^ traces of the models, ALPHAFOLD2 server was used to search for all-atom models. Comparison with experimental data showed all predicted models disagreed equally, as individuals. Finally, we employed molecular dynamics simulations and normal mode analysis-based search coupled with SAXS data to seek better agreeing models. Overall, our analysis concludes that a shifting equilibrium of curved models with low α-helical content best-represents non-associating monomeric α-synuclein.

## Introduction

α-Synuclein (α-Syn) is an intrinsically disordered protein predominantly expressed in various regions of brain. The aggregation of the protein into amyloid fibrils in neuronal cells is associated with cellular toxicity which leads to loss of dopaminergic cells, and onset of the Parkinson’s disease (PD) symptoms. PD is the second most prevalent neurodegenerative disorder which primarily affect motor coordination leading to various symptoms such as akinesia, bradykinesia and tremors (Mhyre, Boyd, Hamill & Maguire-Zeiss 2012). Currently there is no cure for PD and medication only provides symptomatic relief. The α-syn was first discovered in presynaptic terminals and nuclei of neuronal cells (Alderson & Markley 2013). It is a 140 amino acid long protein consisting of an N-terminal α-helix (1-60 residues), highly amyloidogenic non-amyloid-β component (NAC) central region (61–95 residues), and C-terminal acidic tail (96-140residues) (Venda, Cragg, Buchman & Wade-Martins 2010). The N-terminal region contains six to seven 11-amino acid repeats of partially conserved KTKEGV motifs that is involved in interaction with the lipid bilayers (Liu et al. 2021). It is pertinent to mention that most of the known PD-associated pathogenic mutations are located in the N-terminal region of the protein. Interestingly, monomeric α-syn is primarily disordered though its interaction with lipids is known to induce helicity at N-terminal region. Simulations study predict the region 37-45 as the membrane contact region (Tsigelny et al. 2015).The NAC region is primarily responsible for the formation of amyloid-specific cross β-sheet structure. The NAC region is also believed to facilitate α-synuclein association cellular membranes (Anderson, Hirpa, Zheng, Banerjee & Gunawardena 2019). The C-terminal region (96-140 residues) is highly acidic and possesses a random coil structure due to its low hydrophobicity and high net negative charge. This segment plays an important role by protecting the protein from aggregation and degradation (Venda et al. 2010). It is not the loss of protein function that is toxic to cells as mice that lack α-syn exhibit usual behavior and normal dopaminergic cells (Abeliovich et al. 2000). Insight into the preferred conformations of α-syn molecules at different points along its varied pathophysical paths should provide more comprehension in this area.

Different studies with varied techniques have summarized that *in-vitro* purified recombinant α-syn exists predominantly in intrinsically disordered monomeric state which adopts various conformations in equilibrium (Fauvet et al. 2012; Killinger, Melki, Brundin & Kordower 2019). Studies have also shown that the protein isolated under non-denaturing conditions from patient brain tissues or neuronal cells exist as tetramer (Bartels, Choi & Selkoe 2011). It is believed that destabilization of tetrameric form into monomer promotes α-syn aggregation. The fibrillation of α-syn begins with conformational transition of the monomers into aggregation-competent species that further interact to form oligomers (Narkiewicz, Giachin & Legname 2014). The oligomers form cross β-sheet rich ordered assemblies that further grow into fibrils. Monomer binds to the fibrillary end which increases the fibrillary length and mass without increase in their numbers. Studies show that monomer first transiently interact with fibrils using electrostatic interactions, which leads to conformational changes in the α-syn exposing the central NAC region and initiation of secondary nucleation process. The helical propensity differs among α-syn mutants such as its reduced in A30P mutant, but A53T mutant shows no significant changes as compared to the wild type molecule (Tsigelny et al. 2015). Additionally, paramagnetic relaxation enhancement study using spin labelled α-syn and NMR data at 15°C supported that monomeric α-syn exist in multiple conformations which interconvert between nanosecond to microseconds timescales (Bertoncini et al. 2005; Heise et al. 2005).

Among various α-synuclein species, oligomeric intermediates are known to be the most toxic species (Tetzlaff et al. 2008; Vaikath et al. 2022). The structure-activity relation of oligomers has been difficult to discern due to their metastability and transient nature, and also due significant variation in the association state of different species (Vaikath et al. 2022). Studies show that amyloid fibrils are highly polymorphic, and conditions such as buffers, pH, and temperature leads to high variability in fibrillar structure organization (Mehra, Gadhe, Bera, Sawner & Maji 2021; Ziaunys, Sakalauskas, Mikalauskaite & Smirnovas 2021). Similar structural variation has also been observed in samples obtained from human brain extracts which could be the basis of disease symptom variations in synucleinopathies. CryoEM study showed that mature fibrils under cryo conditions form a Greek-key β-sheet topology with parallel β-sheet hydrogen bonds and stabilized by salt-bridges with a hydrophobic core formed by aromatic residues (Li et al. 2018). Further CryoEM studies on wild type α-syn and mutant fibrils proposed existence of two polymorphic species: rod and twister which differ with respect to inter-protofilament interface (Chakraborty & Chattopadhyay 2019; Zhao et al. 2020; Sun et al. 2021; Sun et al. 2023). Additionally, α-syn been coined as a lipophilic protein, its lipid interaction is known to promote its fibrillation and also has an effect on fibrillar structure and cellular toxicity (Fusco et al. 2017; Galvagnion et al. 2019; Sarchione, Marchand, Taymans & Chartier-Harlin 2021). The phospholipid interacts and fills the central cavities present in the fibrils as well as shields the exposed hydrophobic residues significantly influencing the structural organization of protofilament (Galvagnion et al. 2019; Dasari et al. 2022; Dou & Kurouski 2022; Frieg et al. 2022). Not limited by experimental limitations, molecular dynamics simulations have been attempted to decipher structural properties of monomeric and dimeric state of α-syn molecules, and some results have been correlated with independently acquired experimental data (Yu, Han, Ma & Schulten 2015; Zhang et al. 2022; Zamel et al. 2023). It can be safely summarized that in pre-fibrillar monomeric state, a range of interconvertible conformations are accessible and studies to date are constrained by limitations of the experimental technique(s) employed. Since, some conformations are fibril forming competent and fraction of such conformations possibly vary in wild type vs. mutants, and experimental or physiological conditions, more attempts are required to understand of the monomeric state of α-syn. Solution SAXS is a technique, where time- and rotation-averaged data on weighted ensemble of conformations accessible to protein in solution can be reliably acquired (Ashish et al. 2007; Pandey et al. 2014; Badmalia et al. 2018; Ashish 2022). SAXS based shape restoration of the scattering particles is reliable for globular and monodisperse systems, and can be correlated with atomistic structures or models (Sharma et al. 2020; Singh et al. 2020; Chauhan et al. 2023). In contrast, for systems with Gaussian-chain like scattering shape and/or representing transition between states, the dummy residue based models represent weighted average of all conformations accessible to scattering particles at that stage (Ashish et al. 2007; Pandey et al. 2014). As always, newer analysis protocols can be applied to derive new insights in this dynamic protein.

Thus, not limited by the need for diffraction quality crystal or spin-labelled protein in monodisperse conformational state for NMR studies and/or since monomeric state being too small and mobile to be deciphered by CryoEM maps, a number of SAXS studies have been attempted on α-syn protein to understand different stages of fibrillation in solution (Tashiro et al. 2008; Rekas et al. 2010; Ullman, Fisher & Stultz 2011; Curtain et al. 2015; van Maarschalkerweerd et al. 2015; Herranz-Trillo et al. 2017; Chung et al. 2019; Moretti et al. 2020; Ahmed et al. 2021; Lindsay, Mansbach, Gnanakaran & Shen 2021; Panuganti & Roy 2022). Initial studies coupled NMR data with SAXS data to understand range of conformations accessible to α-syn monomer in solution. Importantly, they addressed structural plasticity in this protein (Ullman et al. 2011), and time-dependent SAXS studies which implied that non-β component drive the fibril formation (Tashiro et al. 2008). Another study using size exclusion chromatography coupled with SAXS optics (SEC-SAXS) tracked wild type and some mutants capable of forming fibrils at relatively faster rate (Curtain et al. 2015). The SAXS data profiles from monomeric states of the proteins were analysed using ensemble optimization method (EOM). Wild type α-syn showed prevalence of two major conformer ensembles: compact and extended. Latter extended population was seen to be increased in rapidly fibril-forming mutants of α-syn, E46K and A53T. Another interesting study which tracked formation of fibrils by E46K α-syn by time dependent SAXS data addressed the resident species in samples by Chemometric Data Decomposition (Herranz-Trillo et al. 2017). This study was not limited by assumption of monodispersity in different stages of experiment, and decomposed information by using multiple curves. Overall, key known information from SAXS data analysis of the monomeric stage of α-syn was that this protein molecules adopt Gaussian Chain like or inherently disordered protein (IDP) like solution shape. Some molecular dynamics (MD) simulations in synchronization with SAXS data analysis were also reported which addressed conformational heterogeneity of IDP state of α-syn molecules (Yu et al. 2015; Ahmed et al. 2021). Interestingly, application of meta-interference and Bayesian entropy-based weighing methods with force-field comparisons led to more compact ensembles than seen from SAXS data alone. Considering the advent of new modelling approaches, we revisited α-syn conformations by merging SAXS data from monomeric state with EOM, ALPHAFOLD2, MD simulations and normal mode analysis. Our results do concur with previous findings but goes beyond to provide fresh set of atomistic models which can be used by other interested researchers.

## Materials and Methods

### Protein Expression and Purification

α-Syn expression and purification was performed using method described before with slight modifications (van Raaij, Segers-Nolten & Subramaniam 2006). Briefly, cells were lysed by sonication and further boiled at 95°C for 30 min. Resulting supernatant contained soluble α-syn which was processed for further purification. Streptomycin sulfate and glacial acetic acid precipitated nucleic acid material was removed by centrifugation at 13000 x g for 30 minutes followed by selective α-syn precipitation by ammonium sulphate. The protein pellet was separated, washed with equal volume of 100 mM ammonium acetate followed by equal volume of ethanol and dried for any residual ethanol evaporation. The dried protein pellet was dissolved in buffer containing 10 mM HEPES, 50 mM NaCl, (pH 7.4), dialyzed to remove any residual salts. Protein purity was analyzed using 15% SDS-PAGE. For biophysical experiments, the purified protein was subjected to gel filtration chromatography kept at 10°C (using ÄKTA pure Chromatography System, USA).

### Size Exclusion Chromatography

Freshly purified protein was loaded to Superdex 200 Increase 10/300GL column with flow rate 0.3 mL/min at 4°C. The molecular weight of eluted protein fraction was estimated using elution profile of reference proteins of known molecular weight (Gel Filtration Calibration Kit LMW, Cytiva, GE28-4038-41) (Solanki et al. 2014). Gel filtration chromatography data and standard plot from known reference proteins are mentioned in **Supplementary Fig S2A and S2B**, respectively. Throughout the study, protein concentration was calculated with absorbance value at 280 nm using an extinction coefficient of 5960 M^−1^ cm^−1^ (Ahmed et al. 2021).

### Fibrillation Experiments

Freshly eluted protein was centrifuged at 13000 x g for 30 minutes to remove any insoluble species and used for monitoring *in-vitro* fibrillation. Supernatant with α-syn concentrations in the range of 6-8 mg/mL (and lysozyme as negative control) were incubated at 37°C or 25°C for the indicated time in presence of thioflavin-T and 0.01% sodium azide with continuous linear shaking at 861 rpm (Infinite M plex, TECAN, Switzerland). Fluorescence intensity was recorded at every 15 minutes interval with excitation and emission wavelength of 442 and 485 nm, respectively. The data was plotted against time to visualize the respective change in relative fluorescence units (RFU). For monitoring amyloidic association or seeding mediated fluorescence changes without shaking conditions, 6-8 mg/mL prefibrillar α-syn and lysozyme were incubated in presence of thioflavin-T for 5 hours at 10°C with no agitation. All the data was plotted using Sigma Plot 10.0 software.

### Circular Dichroism (CD) Experiments

Far-UV CD spectra data was analyzed to estimate characterizable secondary structure content in α-syn using JASCO J-815 CD spectrometer. Briefly, a sample of α-syn with concentration of ∼12.5 µM (∼0.18 mg/ml) was used for the CD experiments at 25°C with scanning speed of 50 nm/min at 1.00 nm bandwidth and 0.1 nm data pitch. The recorded CD spectra ranged from 190-250 nm with three total accumulations. The buffer spectra were acquired under the same conditions and averaged profile was subtracted from average for α-syn solution. The data was plotted with the respective mean residue ellipticity (MRE) values using Sigma Plot 10.0 software. Using different databases (as mentioned in the results section) secondary structural content in the protein molecules under the experimental conditions were estimated.

### SAXS Data Acquisition and Processing

Freshly purified α-syn protein was stored in 10 mM HEPES, 50 mM NaCl, pH 7.4 at 4°C and utilized for SAXS experiments within 4 hours of SEC based purification and concentration of protein. All SAXS experiments on α-syn samples were performed at in-house SAXS instrument (SAXSpace, Anton Paar Austria). Line collimation profile of source X-rays were utilized, and scattering information was captured on 1D Mythen detector (Dectris, Switzerland). Optical alignment and data acquisition was controlled by SAXSDrive program. All samples and matched buffers were exposed to X-rays for three frames of 30 minutes at 10°C in the same thermostated quartz capillary and averaged. Three concentrations of α-syn solutions were used for data collection, i.e. stock of 6-8 mg/ml and its two weight-over-weight half dilutions (∼3-4 and ∼2 mg/ml). Acquired datasets were corrected for beam position using SAXSTreat program and further processed for buffer subtraction and desmearing using SAXSQuant program as reported before (Kumar et al. 2021; Vasudeva, Sidhu, Kalidas, Ashish & Pinnaka 2021). We performed direct desmearing of the datasets on buffer subtracted files using line profile of the beam using SAXSQuant program [detailed in the supplementary section of (Goel et al. 2022)]. Resultant datasets represented profiles collected from true point source. No further processing of data was done. The processing provided SAXS profiles of the α-syn molecules in solution at three concentrations mentioned. Since signal-to-noise was significant in the dataset from the most diluted sample, we presented only two concentrations here *i.e*. for 6-8 and 3-4 mg/ml solution. Datasets were in the format of Intensity of scattering, I(s) as a function of momentum transfer vector, s which was defined as 4π(sinθ)/λ with units in nm^-1^. The final column in the datasets were error in I(s) values. All datasets, relevant analysis and models are available at SASBDB database under project IDs SASDQJ7 and SASDQK7.

### SAXS Data Analysis and Dummy Residue Modeling

Particle size parameters like radius of gyration (R_g_) and maximum linear dimension (D_max_) values were estimated using the Guinier approximation of the datasets using in-built plugin in PRIMUS suite of programs version 3.0.1. Kratky plots were also made by same suite to assess globular or Gaussian-chain like behavior of scattering species in solution. The D_max_ values were estimated from the Indirect Fourier transformation of the dataset using GNOM program. This provided a histogram of probability of different interatomic vectors inside the scattering shape of protein molecules, P(r) as a function of vector length in the real space, r in nm. It is to be noted that all further modeling and SAXS profile comparisons were done using the SAXS dataset for 6-8 mg/ml sample. Using the P(r) curve and dummy residues of asparatic acid, ten independent chain-ensemble models were computed using GASBOR program (online version 2.3i r14636). Since there are 140 amino acids in the primary structure of α-syn, for each run, 140 dummy residues and 90 water molecules were used to model shape of α-syn molecule. No further symmetry or shape bias were used during shape restoration. Ten models thus solved were clustered and averaged using DAMCLUST program (ATSAS suite 3.0) (Manalastas-Cantos et al. 2021). Briefly, the routine employed SUPCOMB program (Kozin & Svergun 2001) to align inertial axes of ten models and clustered models based on their closeness in spatial disposition. Similar structural models were aligned and averaged using DAMAVER program (Volkov & Svergun 2003) in the DAMCLUST routine (Petoukhov et al. 2012).

### SAXS Data based Ensemble Optimization Method (EOM)

Using online version of EOM v 3.0 (Tria, Mertens, Kachala & Svergun 2015), ensemble of conformations accessible to α-syn molecules in solution were solved (Curtain et al. 2015; Badmalia, Singh, Garg & Ashish 2017). It is a suite of programs which uses a pool of independent models available in structural database. Pool generation was done using genetic algorithm (GAJOE) program version 2.1 - (r14636). During selection of pool or ensemble of structures to search, flexibility of the system under study was kept into consideration. This was assessed from the Kratky plot of the dataset or orthogonal information available about the protein. For Gaussian-chain like system, a wider distribution was considered to reflect its disorderly behavior in solution. Flexibility of the pool was computed using ratio of theoretical Rflex vs. Rsigma. Rflex is 100% and 0% for fully flexible and fully rigid system, respectively, and Rsigma approaches 1.0 for fully flexible system and less than 1.0 for systems with significant flexibility. Sequence information of the α-syn protein, experimental SAXS profile, and random coil nature of the protein (as seen from CD data and Kratky analysis) were inputs to search for suitable templates. In the present study, two runs of EOM were done: 1) considering, all 140 residues to be disordered, and 2) first 60 residues were presumed to be α-helical. For intrinsically disordered or unfolded portions, no templates of rigid or globular bodies were considered, and random configurations of the C^α^ trace models were generated using RANCH (RANdom CHains) [ATSAS 3.1.3 (r14636)] program in EOM suite of programs. For each run, 10000 theoretical curves from templates were searched to obtain best-fits. 20 curves were considered per ensemble and 100 cycles of genetic algorithm during search process. RANCH program provided *.cif format of the solutions, and their respective R_g_, D_max_ and fraction representation in experimental data. These coordinate files were resaved in PDB format, as the CIF format was found to be incompatible with used version of CRYSOL program (both off- and online) (Svergun, Barberato & Koch 1995).

### Predictions from AlphaFold2 server and Comparison with SAXS data

Using the primary structure of α-syn as used for SAXS experiments (**Supplementary Figure S1**), ALPHAFOLD2 server was accessed at https://colab.research.google.com/github/sokrypton/ColabFold/blob/main/AlphaFold2.ipynb (Jumper et al. 2021). Protocol employed multiple sequence alignment with available templates and accessible databases. Using default parameters, and searching for monomeric version of protein, lDDT (Local Distance Difference Test) was predicted at each residue level of queried sequence (Mariani, Biasini, Barbato & Schwede 2013). Accordingly, five models were finalized which were ranked as per pan-sequence predicted lDDT values. Models were relaxed using AMBER force field under default conditions, and for each 3D model, Predicted Aligned Error (PAE) was generated (Guo et al. 2022). Further, a plot indicating number of sequences identical to resembling query was provided as a function of residue position. CRYSOL program (Svergun et al. 1995) was used to compute SAXS profiles of the top ranked models and compare with the experimental SAXS data. For each pair of theoretical vs. experimental profiles, χ^2^ value was computed as a quantitative measure of similarity. χ^2^ value of unity is indicator of identical profiles, and deviation from value of 1 (both less or more) indicates disagreement between the respective two datasets. PyMOL program was used to visualize PDB models, alignments and generation of figures.

### Molecular Dynamics Simulations (MDS)

All MDS runs in this were done using YASARA Program v6 (Krieger, Koraimann & Vriend 2002; Krieger & Vriend 2014; Ozvoldik, Stockner & Krieger 2023) and a computing system having 16 core processors plus four GPUs. The five models predicted from ALPHAFOLD2 server were subjected to MDS individually. YAMBER2 forcefield was employed. The protein structures were added with all hydrogen atoms, soaked in cuboidal TIP3P water system with 5 Å additional space in all three axes, and neutralized with 0.9 % NaCl at pH 7.4. Periodic boundary conditions were imposed, and non-bonded calculations were cut-off at 8 Å. The solution boxes were energy minimized to a cut-off gradient of 0.01, and then coupled to heating at 283 K for MD runs. Each simulation was run for 10 ns with restart coordinates written out at 2.5 fs. Post-runs, macros were employed to analyze molecular and simulation cell features. For this work, primary focus was on the percentages of secondary structural content, per residue structure, theoretical R_g_ values and per residue Root Mean Square Fluctuations (RMSF) values as a function of MDS time. Visual analysis of the resultant PDB files was done by PyMOL program.

### Normal Mode Analysis Based Search for Better-fitting Models

Taking top five ranked models solved from ALPHAFOLD2, SREFLEX program was employed to search for close structures accessible to starting structure within low frequency normal modes of vibration and agreeing to experimental SAXS data (Panjkovich & Svergun 2016). ATSAS 3.0.0 (r11734) offline version of the program was used and the first 15 modes were explored. The program’s algorithm searched in hierarchical order, by first considering large scale rearrangements in the starting PDB file, and then considering smaller/localized changes. At all steps, improvement in the agreement between theoretical SAXS profile of the model vs. experimental data was monitored and used as logic for further search. Normal mode analysis considered C^α^ atoms in the PDB structures of α-syn as centroids for perturbation and follow-up. For each search, both restrained and unrestrained models were computed with nomenclature of rc*.pdb and uc*.pdb, respectively in PDB format. Here, restrained models gave higher weightage on retention of backbone stereo geometry vs. more allowed of deviation in the unrestrained models. Furthermore, for each solution, χ^2^ between theoretical vs. experimental SAXS profiles, RMSD between centroids of initial and resultant structure, breaks and clashes in solved model were written out along with hierarchy of the model. As before, the models were visually assessed using PyMOL program and the same was used to make figures.

## Results and Discussion

### Protein Purification and Characterization

α-Syn was purified as described in the Materials and Methods section. The theoretical molecular weight of monomeric α-syn is 14.46 kDa. The Gel-filtration experiments were performed at 4°C to estimate the hydrodynamic mass of the protein using protocol similar to as described before (Solanki et al. 2014). The α-syn was consistently obtained as one single peak at elution volume higher than that expected for its monomeric molecular weight (**Figure S2A** and **S2B**). The α-syn elution peak was estimated to be between elution volume corresponding to monomeric conalbumin (75 kDa) and ovalbumin (44 kDa) which is consistent with an earlier study (Fauvet et al. 2012). Increased elution volume value than expected from globular protein of similar mass could attributed to its non-globular or unfolded elongated structure (Paleologou et al. 2008). The fraction eluted from gel filtration column was further loaded onto Native-PAGE. Only single band appeared on the Native-PAGE suggesting presence of single oligomeric species in solution. In agreement with the previous studies (Fauvet et al. 2012; Araki et al. 2016), our results also indicated that our gel filtration chromatography purified fractions are predominantly monomeric α-syn which was used for further experiments.

### Monitoring time and temperature dependent Thio-T mediated fibrillation

Fibril formation by freshly gel filtration purified α-syn at final concentration of ∼6-8 mg/ml was confirmed using thioflavin-T dependent kinetics. As expected, at 37°C under rigorous shaking condition, with a lag time of about 2 hours, the protein showed significant fibrillation with distinct primary nucleation, followed by a secondary nucleation and saturation of fibril formation **(Fig. 1A)**. Importantly, the delay time for initiation of fibrillation were much delayed when α-syn was incubated at 25°C with all other kinetic parameters unchanged **(Supplementary Fig. S2D)**. Similar observations have been previously reported as well (Ohgita, Namba, Kono, Shimanouchi & Saito 2022). Our results and prior reports clearly conclude that temperature significantly impacts initiation of fibril formation. It is pertinent to mention here that at lower temperature i.e. at 10°C, no significant fibril formation was observed even after 5-6 hours (**Fig. 1B**). Results were comparable to the negative control used here, i.e. lysozyme. Taking cue from these results, we planned and completed CD and SAXS experiments in this time window, where dismal fibrillation was observed. This ensured that the data was acquired from predominantly monomer/ unassociated population of α-syn.

**Figure 1.**
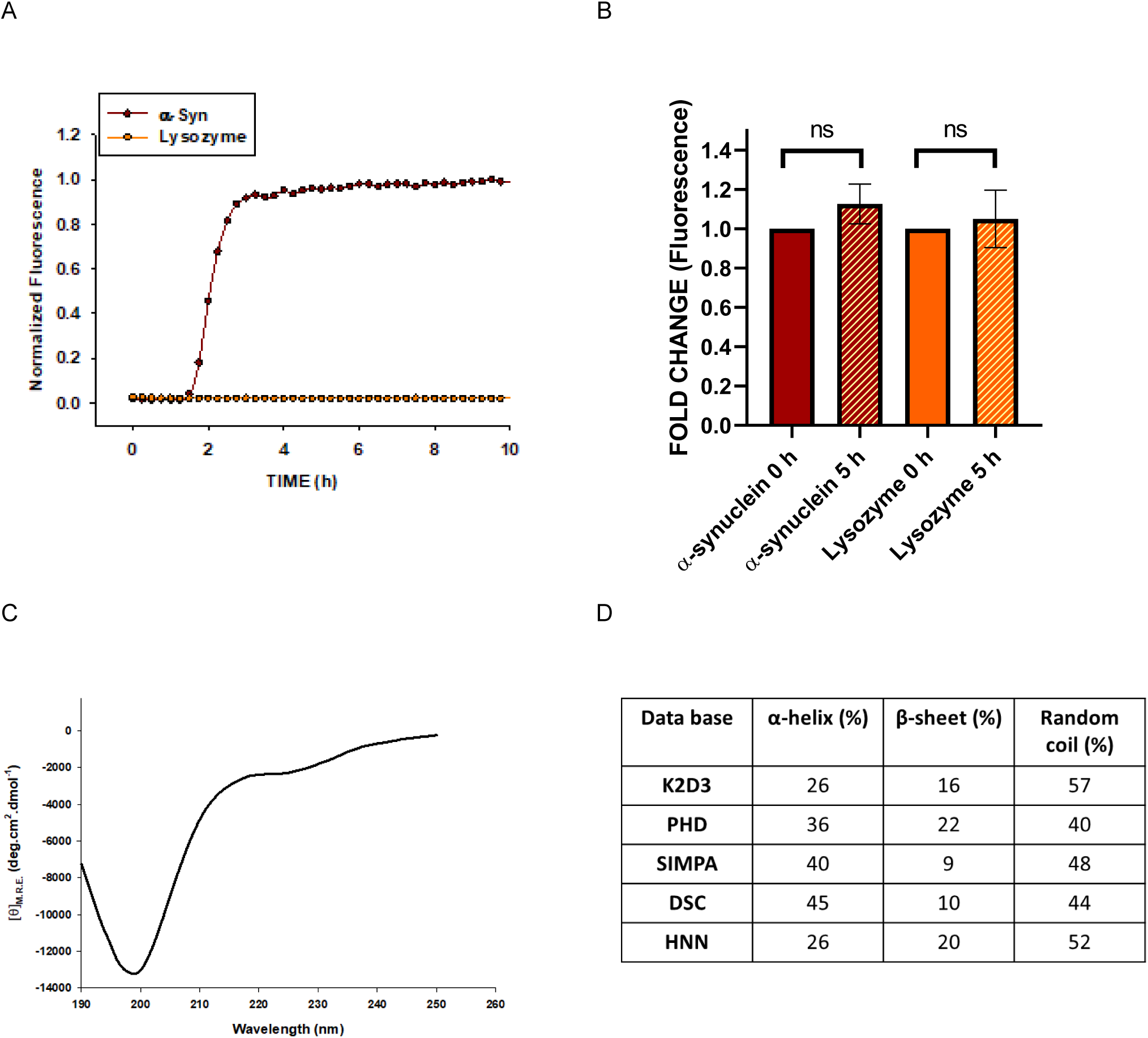
α-Syn *in-vitro* thioflavin-T mediated fluorescence measurements and secondary structural analysis data are presented here. (A) Thio-T mediated fluorescence measurement of pre-fibrillar α-syn at 37°C in shaking condition and (B) thioflavin-T mediated fluorescence kinetics at 10°C with static conditions showing changes in aggregation mediated fluorescent intensity changes. (C) Far-UV CD spectra of prefibrillar α-syn showing their observed secondary structural pattern. (D) Secondary structural content (in percentages) was analyzed from the acquired CD spectra with different prediction servers.

### CD Data Analysis of the Pre-fibrillar state α-syn

At concentrations of 12.5 μM (∼0.18 mg/mL) stored at 10°C, our sample of α-syn was primarily in monomeric non-fibrillar state. It is to be noted that the temperature of CD experiments was 25°C. As mentioned earlier too, low concentration of protein and temperature used in this experiment were not favorable condition for fibril formation (Sode, Ochiai, Kobayashi & Usuzaka 2007). As expected, the CD spectra for this protein showed presence of random coil (Ghosh et al. 2015) **(Fig. 1C)**. Acquired CD data was processed by different databases like K2D3, PHD, SIMPA, DSC and HNN to deconvolute characterizable secondary structural content in our samples (Rost, Sander & Schneider 1994; King, Saqi, Sayle & Sternberg 1997; Lin, Simossis, Taylor & Heringa 2005; Louis-Jeune, Andrade-Navarro & Perez-Iratxeta 2012; Ema, Sultana, Shaj & Galib 2022). Most of them concluded dominantly random coil like structures due to presence of sharp negative peak around 197-200 nm range and slight negative ellipticity around 230 nm **(Fig. 1D)**. Average of all databases suggested: α-helix to be about 35± 8.4%, β-sheet to be about 16± 5.8%, and random coil to be around 48± 6.7%, respectively.

### SAXS Data on the Pre-fibrillar State

SAXS datasets acquired on two concentrations of α-syn protein in the buffer mentioned in methods are shown in **Fig. 2A**. Timing of experiments were planned in a manner that there was insignificant presence of associated species of α-syn. Double log plot confirmed lack of any aggregation or interparticulate affect in the samples (**Fig. 2A right**). Linearity of the Guinier region of the datasets are presented in **Fig. 2B** and the hyperbolic profiles of the corresponding normalized Kratky plots (**Fig. 2C**) confirmed disordered or Gaussian-Chain like scattering profile of the protein molecules in solution. If the molecules adopted globular scattering profile, in normalized Kratky plots, the curves would have adopted peak like profile with maxima at sR_g_ close to 1.73 (please see the dotted lines in plots in **Fig. 3C**). Linear fit to the Guinier regions of the SAXS datasets at two concentrations of α-syn molecules provided R_g_ value of this protein to be about 4.7 nm. P(r) estimation of the two datasets suggested D_max_ of the molecules in solution to be about 17-18 nm (**Fig. 3A**). It is pertinent to highlight here that the relative frequency of interatomic vectors above 14 nm were less in the estimated P(r) curves and had higher uncertainty suggesting significant variation in long interatomic vectors in frequency and relative orientation in scattering shape of molecules. Significant deviation in the observed D_max_ value of 16-18 nm from 2R_g_ (∼10 nm) implied a rod-like shape of the molecules. Please see the SASBDB depositions, SASDQJ7 and SASDQK7 on the α-syn protein at ∼8 and ∼3.8 m/ml, respectively to evaluate the SAXS datasets and primary analysis. Based on intensity values estimated at zero angles (I_0_) for well-characterized proteins, molecular mass of α-syn molecules in solution were estimated to be about 15-17 kDa (vs. expected value of 14 kDa). The primary analysis indicated that prefibrillar α-syn molecules were monomer in solution with average R_g_ and maximum D_max_ of 4.7 and 16-18 nm, respectively.

**Figure 2.**
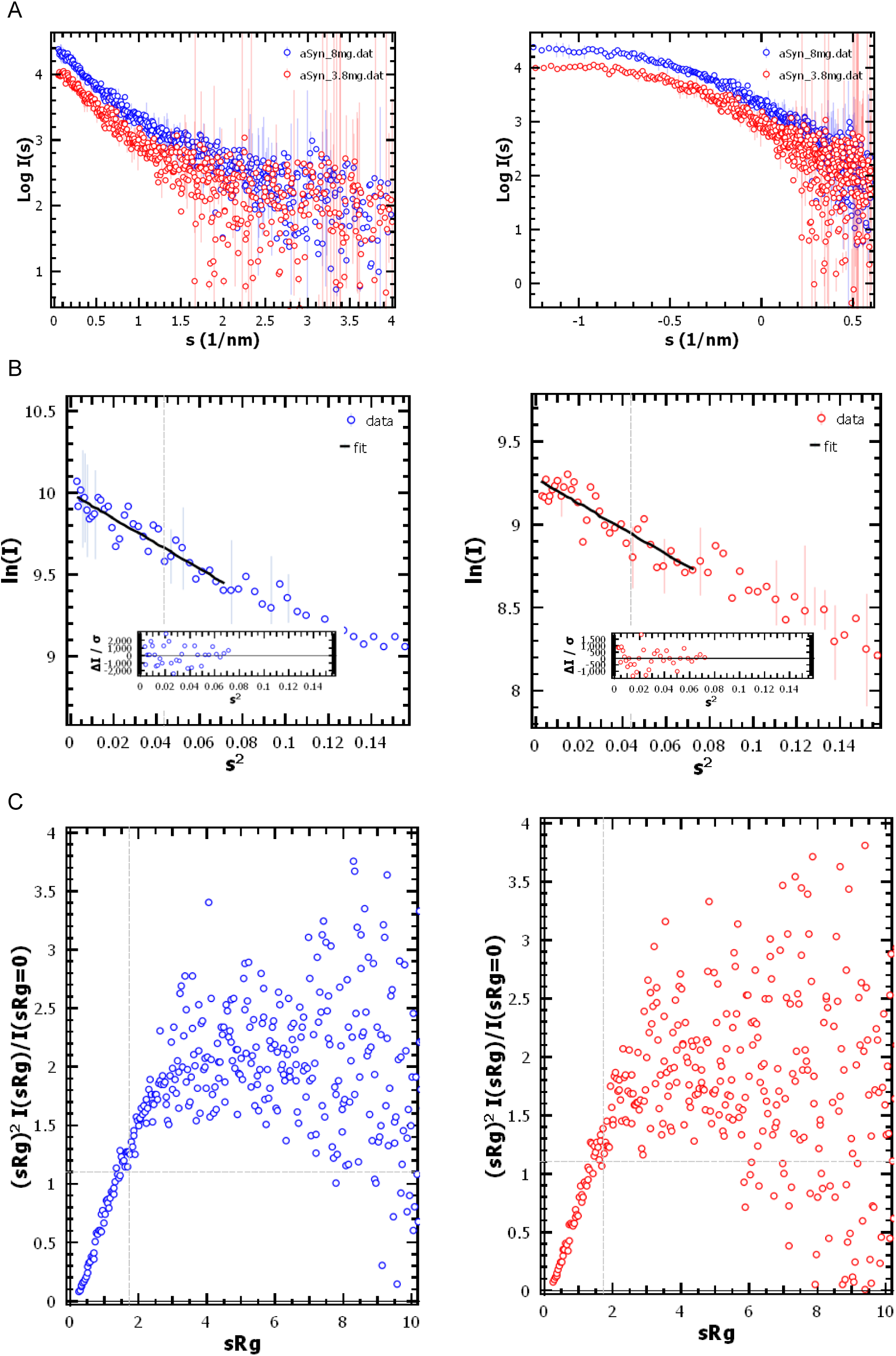
SAXS data acquired on samples of α-syn are presented here. (A) SAXS data from protein concentration of ∼8 mg/ml (blue circles) and ∼3.8 mg/ml (red circles) are shown in two plot formats. (B) Their Guinier fits presuming globular scattering shapes are presented. Black line shows the linear fit and region used to estimate R_g_ values. Insets show the fit residuals. (C) Normalized Kratky plots of the SAXS datasets indicate their Gaussian-chain like scattering profile.

**Figure 3.**
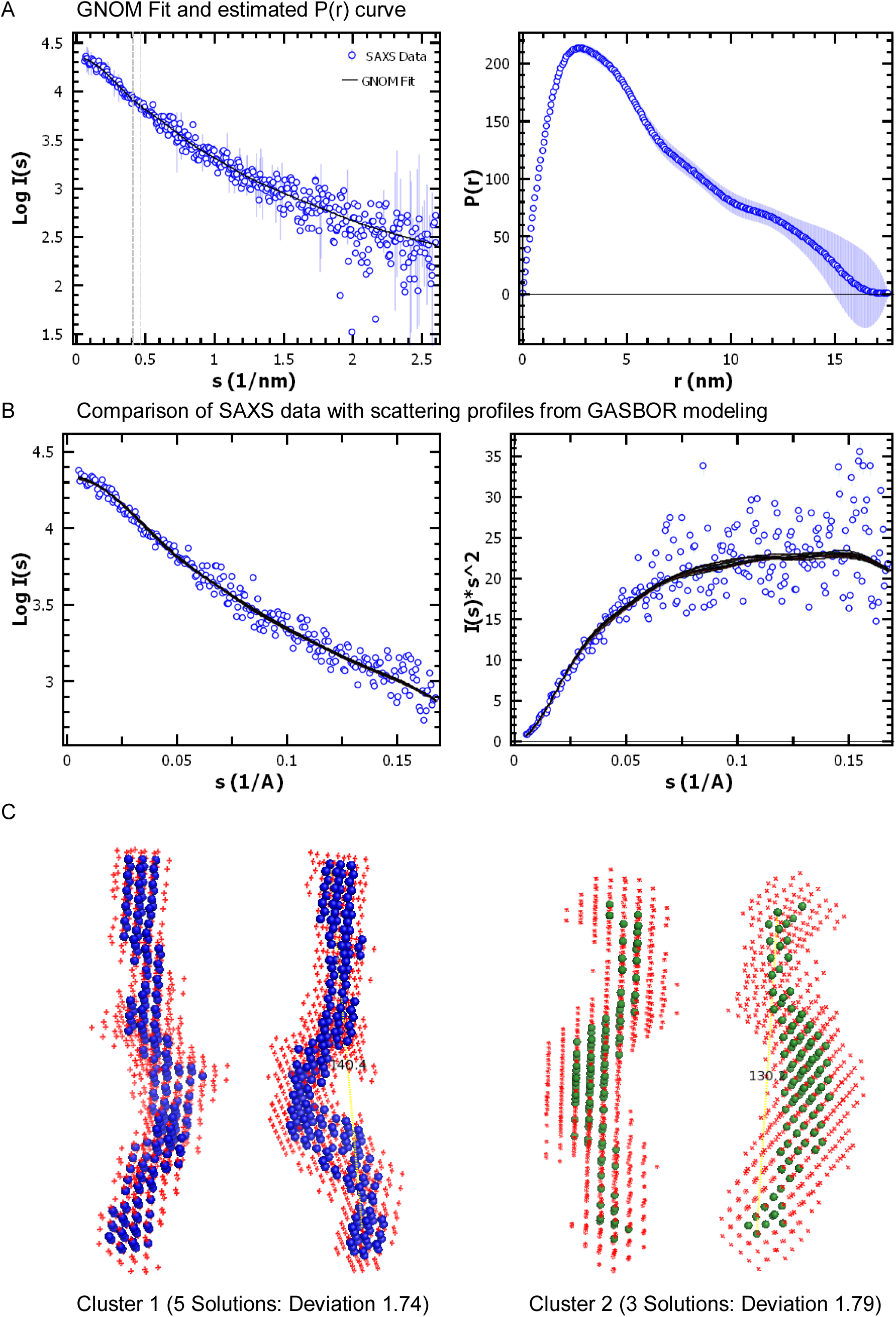
GNOM fit and results of chain-ensemble modeling are shown here. (A) (Left) The black line represents the SAXS profile over the SAXS data (blue circles) which resulted from estimating pair-wise distribution of interatomic vectors, P(r) (shown in right panel). (B) Left panel compares the theoretical SAXS profiles of the ten solutions of Chain-ensemble models of αSyn (black lines) to experimental data (blue circles). Right panel shows the Kratky plot of the comparisons shown in left panel. (C) Two rotated views of averaged models solved for αSyn protein molecules are shown here. These are average of two dominant clusters of solutions. Views are rotated along long axis. The blue and green spheres represent the averaged model and red crosses around depict variation in the models averaged.

Using the estimated SAXS profile and P(r) curve information (deduced from GNOM analysis), chain-ensemble shape restoration runs were done using GASBORIQ program (online version). Ten independent runs were done without any shape or symmetry bias. For each run, 140 dummy residues were considered in-synchronization with that many residues in our protein at monomeric state. In **Fig. 3B**, black lines represent the theoretical SAXS profiles of the dummy residue models solved, and they are compared with experimental SAXS data. Hyperbolic profiles of the Kratky plots in **Fig. 3B** (right) confirmed that the Gaussian-chain like shape of molecules in solution was retained in the chain-ensemble models, solved for α-syn molecules. All ten solutions were clustered as per similarities within their size and shape profiles. Clustering and averaging were done using DAMCLUST and DAMAVER programs, which yielded two main clusters: Cluster 1 with five member solutions and a normalized spatial deviation (NSD) of 1.74 amongst them, and a minor cluster i.e. Cluster 2 with three member solutions and NSD of 1.79. Models and respective deviations are shown in **Fig. 3C** which showed curved spindle shape for the α-syn molecules. Averaged models for Cluster 1 and 2 showed semi-extended shapes with end-to-end dimension of about 140 and 130 Å or 14 or 13 nm, respectively.

### Ensemble Optimization Method (EOM) Based Predictions

Using EOM program, primary structure of α-syn protein and acquired SAXS data, a pool of structural templates was searched to get C^α^ trace models of the protein under study and in agreement with its scattering properties in solution. As mentioned in methods, two runs were done, considering: 1) all 140 residues to be disordered, and 2) first 60 residues to be adopting α-helical structure and rest remaining disordered. For the first run, Rflex (random)/Rsigma was 71.7% (∼83.6%) which meant that the Rflex of the selected ensemble was ∼72% and less flexible than ∼84% for the pool (**Supplementary Fig. S3A**). The templates which supported weighted representation in the experimental SAXS profile were searched and shown vs. the pool (**Supplementary Fig. S3B**). Overall, four models were generated from RANCH program (**Fig. 4**). Comparisons of the computed SAXS profiles of the models with experimental data are shown in **Fig. 4A**. Template number from the pool, χ^2^ value vs. experimental SAXS profiles, and their mole fraction of representation in the selected ensemble are mentioned below the models in **Fig. 4B**. It is to be noted that models 3699, 6492 and 7083 implied 0.32, 0.25 and 0.25 fractions of the overall population of conformations of α-syn as per SAXS data. The R_g_ values were in the range of 32.6 to 46.7 Å. Interestingly, they adopted shapes which resemble the two dominant clusters from chain-ensemble modeling (**Fig. 3C**). The models in **Fig. 4** are colored as per spectrum in primary structure and superimposition of N-terminal 20 residues provided an ensemble where the C-terminal appeared to occupy varied positioning in space. This correlated with variation in certainty values of the long-distance vectors estimated P(r) analysis from experimental SAXS data (**Fig. 3A right**).

**Figure 4.**
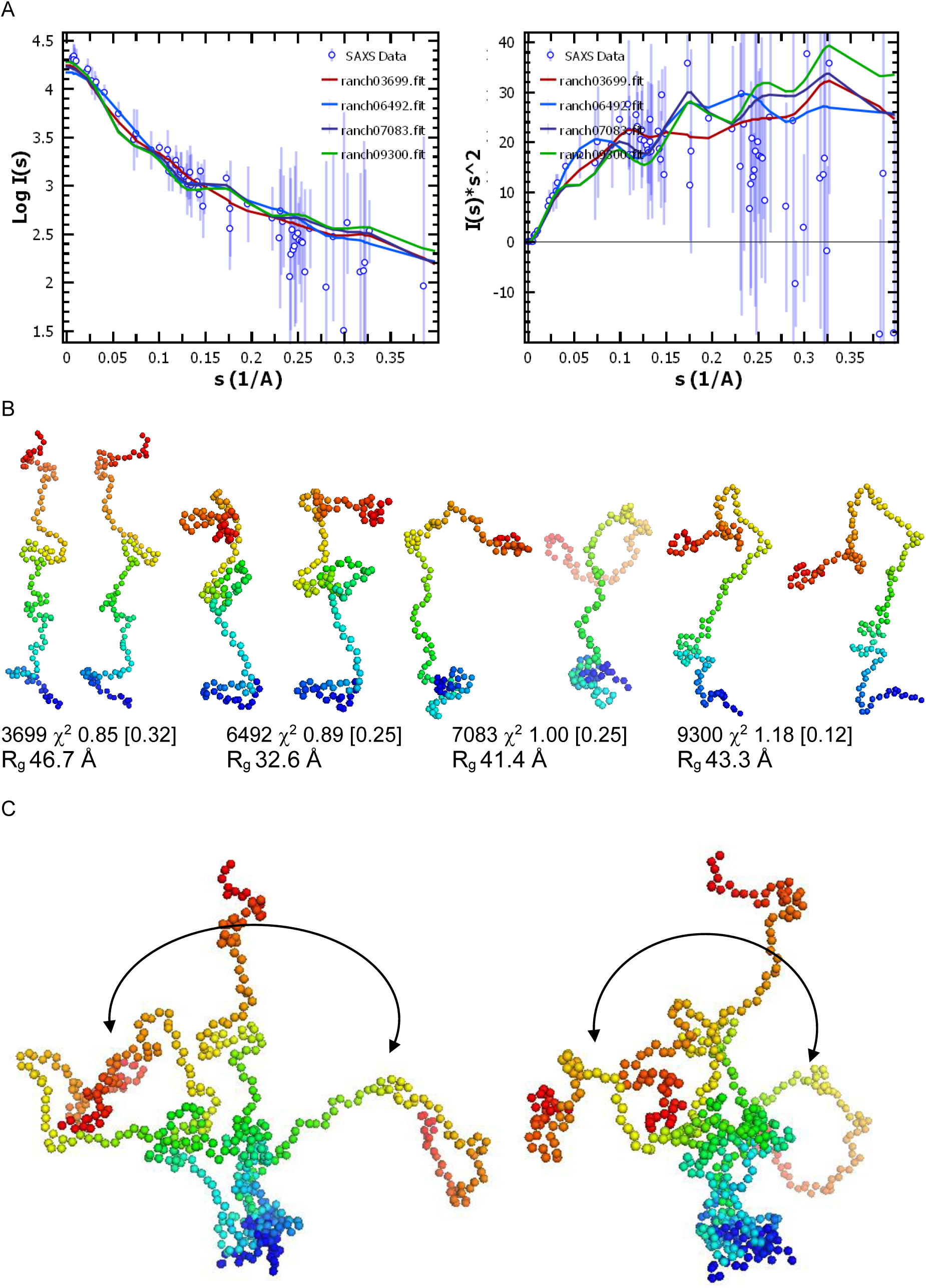
Results from Ensemble Optimization Method (EOM) considering all 140 residues as disordered are displayed here. (A) Left plot shows the comparison of the calculated SAXS profiles of the models searched by EOM (lines) vs. experimental SAXS data (blue circles). Respective model number as solved by EOM are mentioned as legends. Right panel has the Kratky plot of the graph shown on left. (B) Two rotated views of the C^α^ trace models (sphere mode) of the solutions computed by RANCH program are shown here. Respective model number is mentioned below. Coloring has been done using spectrum mode where blue and red represent the first and last amino acid in the primary structure of αSyn protein. Computed χ^2^ value between respective model’s calculated SAXS profile vs. experimental data are mentioned along. Additionally, fraction this model may represent experimental data in ensemble are mentioned in parentheses. (C) Two rotated views of all four solutions aligned across their first 20 residues highlighting variation in the conformations in ensemble.

In the second run, first 60 residues were considered as helix which shifted the overall disorder to last 80 residues in the protein structure. The reason for this assumption is clearer in the follow-up section, where results from another search protocol provided models with high helical content across N-terminal half of protein. For this run, Rflex (random)/Rsigma was 69.8% (∼84.9%) which meant that the Rflex of the selected ensemble was 70% and less flexible than 85% for the pool (**Supplementary Fig. S4A**). It was interesting that despite fixing helical order for the first 60 residues, the flexibility of the system and searched pool were not very different than the first run of EOM. The templates which supported weighted representation in the experimental SAXS profile were searched and shown vs. the pool (**Supplementary Fig. S4B**). Overall, three models were generated from RANCH program (**Fig. 5**). Comparisons of the computed SAXS profiles of the models with experimental data are shown in **Fig. 5A**. From the second run of EOM, selected template number from the pool, χ^2^ value vs. experimental SAXS profiles, and their fraction of representation in the selected ensemble are mentioned in **Fig. 5B**. Models 2712, 7065 and 9085 implied 0.30, 0.20 and 0.50 fractions of population of conformations of α-syn while considering N-terminal to be organized. The Rg values of the models were 28.9 to 44.7 Å (slightly smaller than models solved in above EOM run). As done above, superimposition of the selected templates provided a view that if the first 20 residues are overlapped in space and first 60 residues are helical, then remaining residues stretched out in space like loose chains. This was expected as higher disorder was considered for C-half of protein compared to first EOM run where overall disorder was distributed over the whole sequence. One more relevant observation was that the first EOM run resulted in models with χ^2^ values closer to unity than in the second run. Overall, the two EOM runs supported that the fixation of first 60 residues as helix did not provide better convergence than not having any bias as done in the first run.

**Figure 5.**
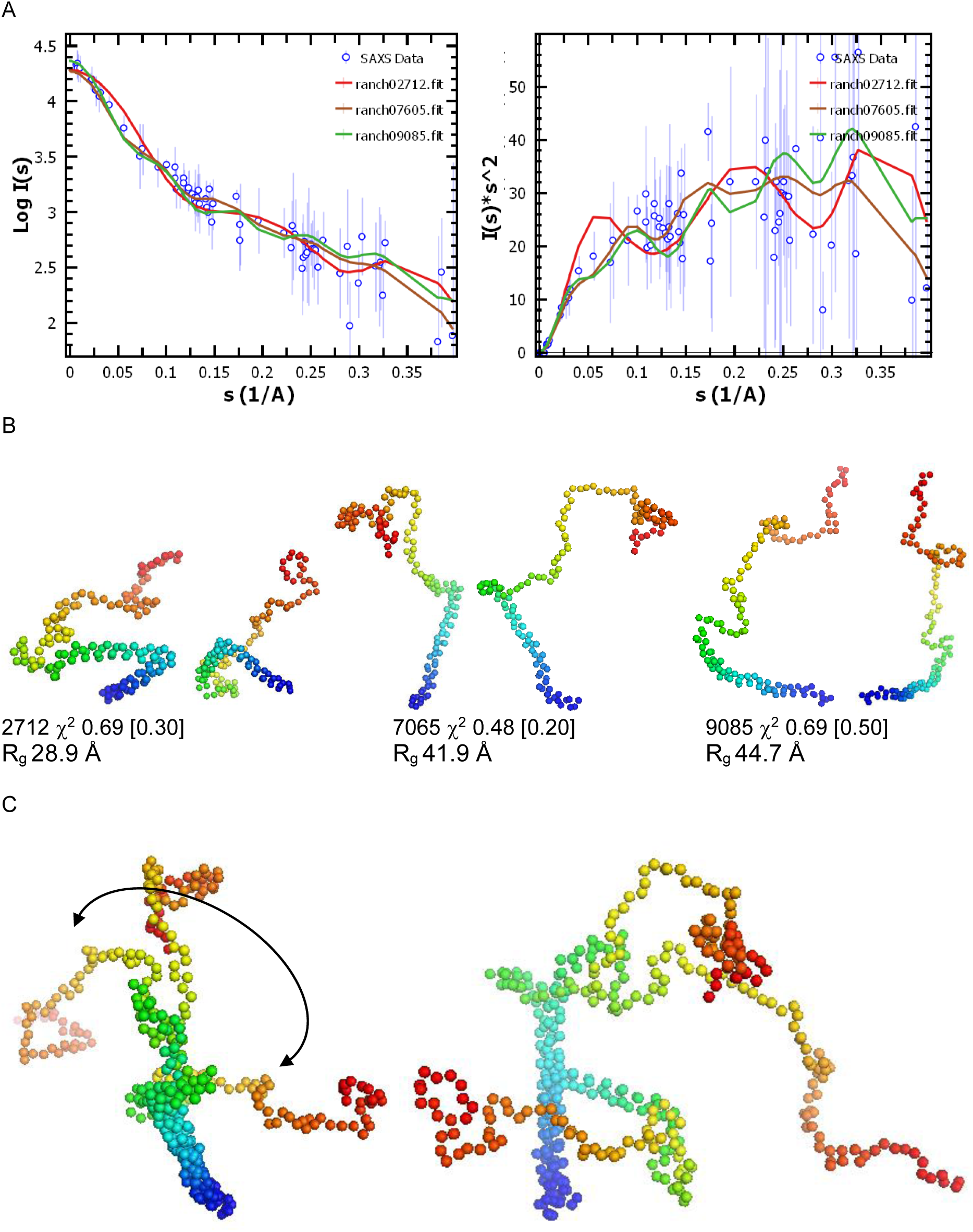
Results from Optimization Method (EOM) considering first 60 residues to be adopting α-helical structure and remaining residues were considered to be disordered, are presented here. All panels have similar description as in respective panels in Figure 4, except that this search provided only three models as solutions.

### Results from ALPHAFOLD2 server

For the sequence of α-syn in the **Supplementary Figure S1**, the ALPHAFOLD2 server provided five top ranked solutions with all-atom structural details (**Figs. 6 and S5**). Theoretical SAXS profiles were compared with experimental data and their χ^2^ values are shown and reported in **Fig. 6A** and **6B**, respectively. Computed R_g_ values of the models are mentioned below each model’s representations which were much smaller than 47 Å, experimentally estimated value. The comparative Kratky plot showed that the disorder present in the experimental data was represented in the models solved. The five top ranked models were in two classes: hairpin shaped helical structures (1^st^ – 3^rd^ ranked models) and curved structures (4^th^ and 5^th^ ranked models). For top 3 and 5^th^ ranked models, there was significant coverage of N-terminal residues in α-helical order, possibly due to use of templates of crystal structures with hydrophobic partners. The χ^2^ values were almost equally deviated from unity suggesting similarity in their probability of existence in the sample. This was also correlated with similar lDDT, PAE and sequence coverage values for the five models (**Fig. S5**). As done before, we superimposed all the models across first 20 residues which provided an impression that α-syn molecules wobble at their C-terminal half if the N-terminal is structured and aligned. This correlated with C^α^ trace models in shape profiles solved using EOM calculations, but these ALPHAFOLD2 server-based residue detailed models were substantially smaller in their R_g_ values than experimental estimations and EOM results. Importantly, only rank 4 model agreed with the lack of secondary structural content as deduced from CD data of this protein.

**Figure 6.**
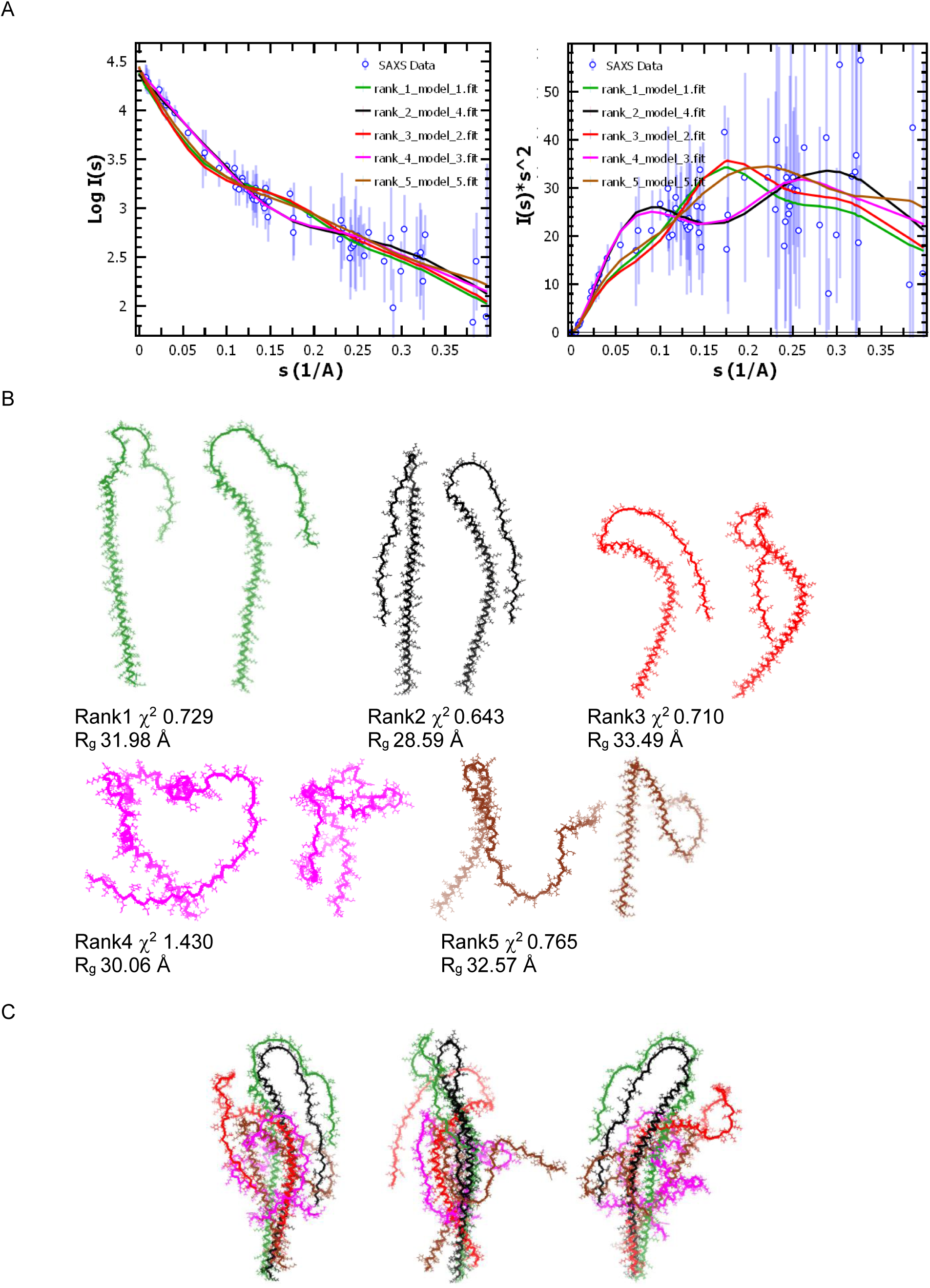
Models solved by AlphaFold2 server and their comparison to experimental SAXS data are shown here. (A) Left plot shows the comparison of the calculated SAXS profiles of the models computed by ALPHAFOLD2 server (lines) vs. experimental SAXS data (blue circles). Respective ranked model number is mentioned as legends. Right panel has the Kratky plot of the SAXS datasets in the left panel. (B) Two rotated views of top five ranked models are shown as ribbon models. Color of the models is same as the color of the line in plots in panels in the A section. Computed χ^2^ values between respective model’s calculated SAXS profile vs. experimental data are mentioned along. (C) Superimposition of the five models are shown after alignment of their first 20 residues.

### MDS Runs of the Top Ranked Models

To explore if the top ranked models from ALPHAFOLD2 server-based predictions can lead to structural models which are extended, and better represent ensemble accessible to α-syn in solution, we performed MDS runs with each individual model. As mentioned in the methods section, each model was input for MDS for 10 ns in explicit water and 0.9 % NaCl at 283 K. Their secondary structural content (total in % and per residue) as a function of simulation time are shown in **Fig. 7** and **Supplementary Figure S6**. This revealed that except Ranked 4 model, all other models retained substantial α-helical structure in their N-terminal stretch. During its MDS run, ranked 4 model preferred secondary structures of α-helix and random coil similar to CD data based interpretations. Analyses also showed that computed R_g_ values increased for all models, but decreased for ranked 4 model which folded in a manner that its two terminals came together. This indicated that during MDS runs while 1-3 and 5^th^ ranked model relaxed open during simulation. Using experimental R_g_ of 47 Å as constraint, very few structures from MDS runs of 1^st^, 2^nd^ and (little more) for 3^rd^ ranked models could be considered to represent information in experimental SAXS profile (shown as open black box in R_g_ variation plots). For MDS runs of 4^th^ and 5^th^ ranked model, we selected only structures which had highest R_g_ values of their trajectories. Comparison of the averaged computed SAXS profiles of the selected structures with experimental data is shown as Kratky plot and χ^2^ values in **Fig. 7**. For 1-3 top ranked model, the computed χ^2^ values suggested that MDS aided in improving starting structure to shift towards SAXS profile. The average of the selected models from MDS trajectories indicated bends in the helical order and a clear shift from the hairpin-like shape predicted for top three ranked by ALPHAFOLD2 server. For structures selected from MDS runs of 4^th^ and 5^th^ model, there was a clear disagreement between two datasets in their Kratky plots (highlighted with red arrows). This was expected as we selected structures which Rg values much smaller than experimental value of 47 Å. In correlation, computed χ^2^ values also indicated that MDS runs did not help for 4^th^ and 5^th^ ranked models, though percentages of helical and random coil content matched best for 4^th^ ranked model.

**Figure 7.**
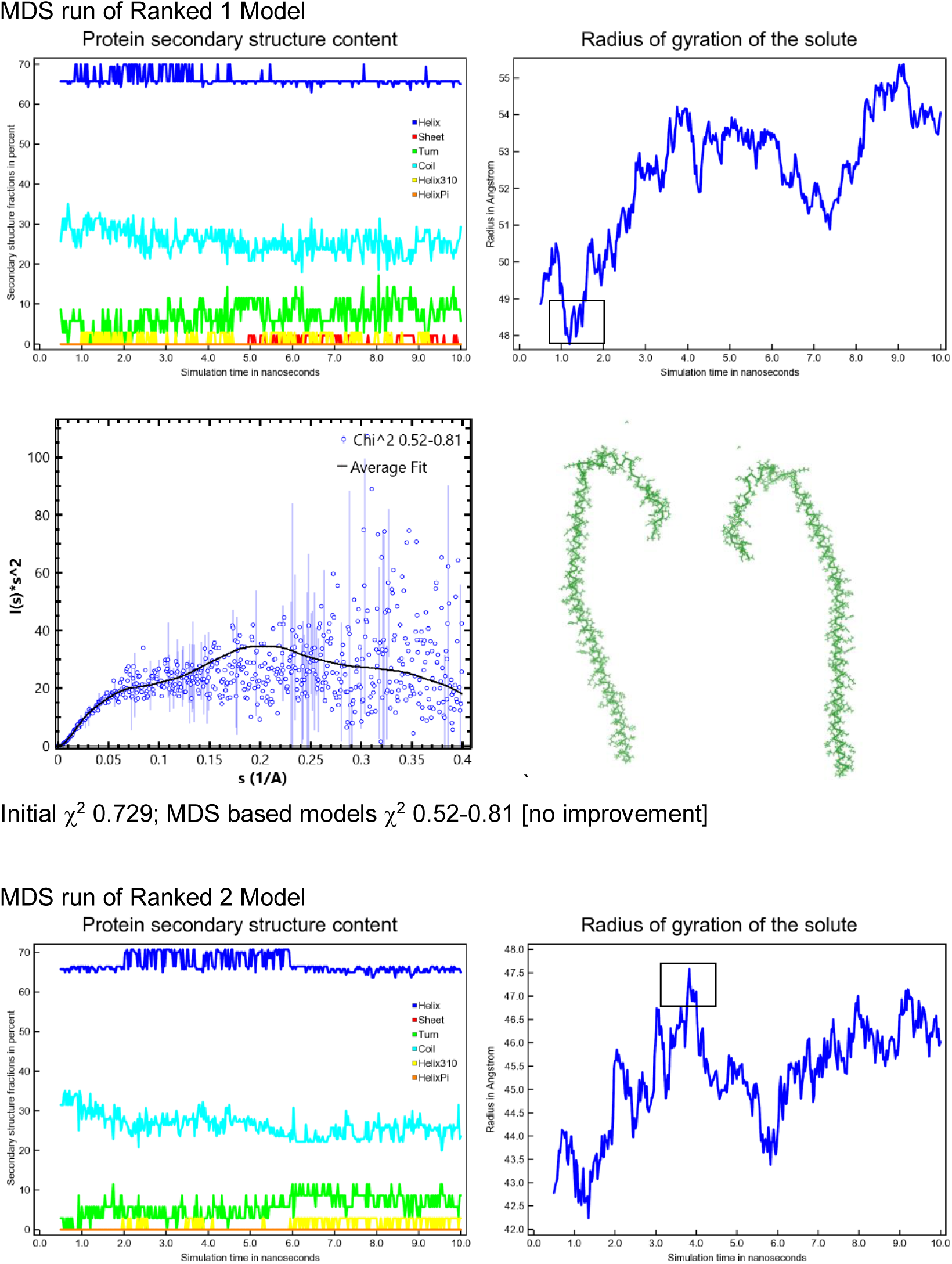
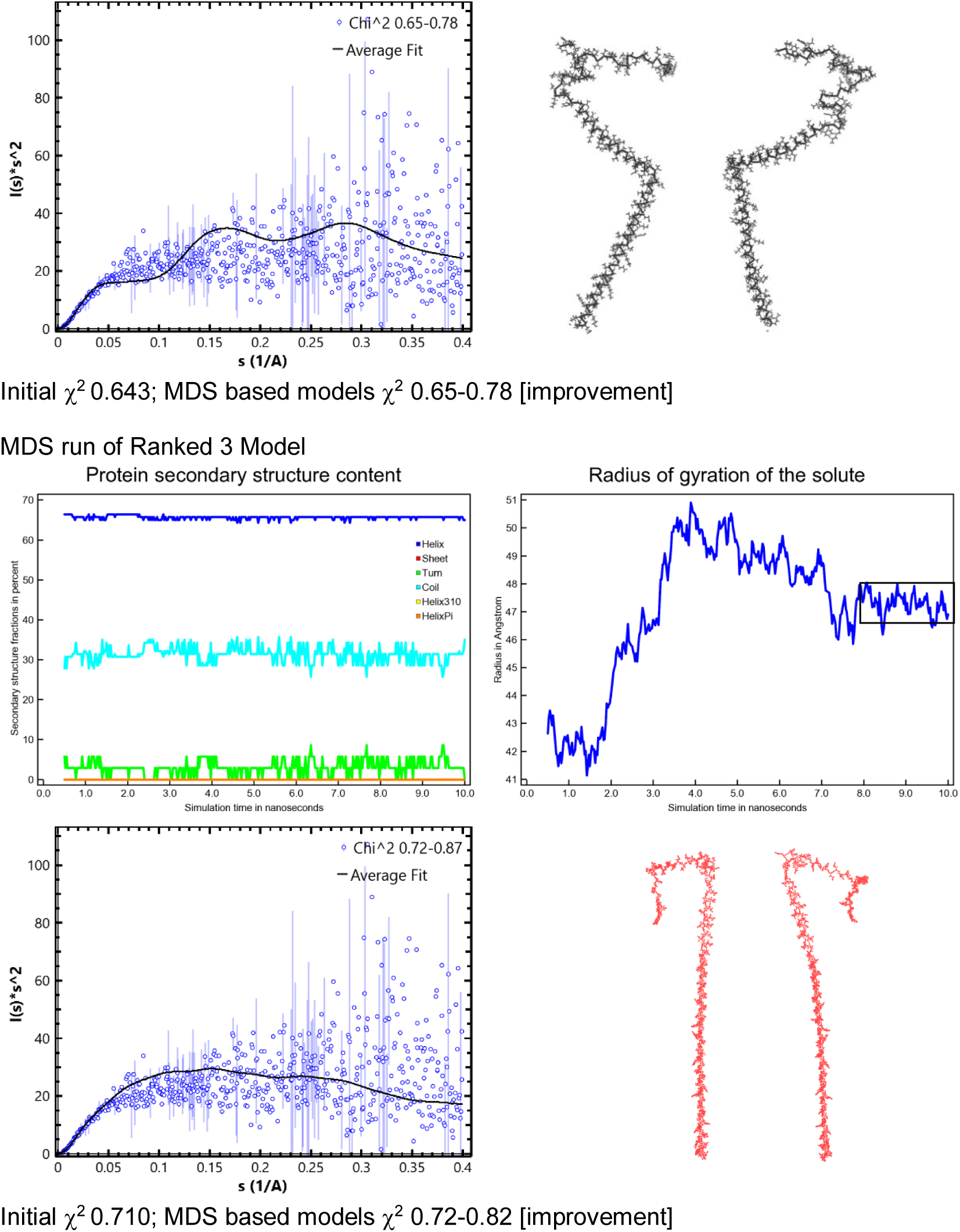
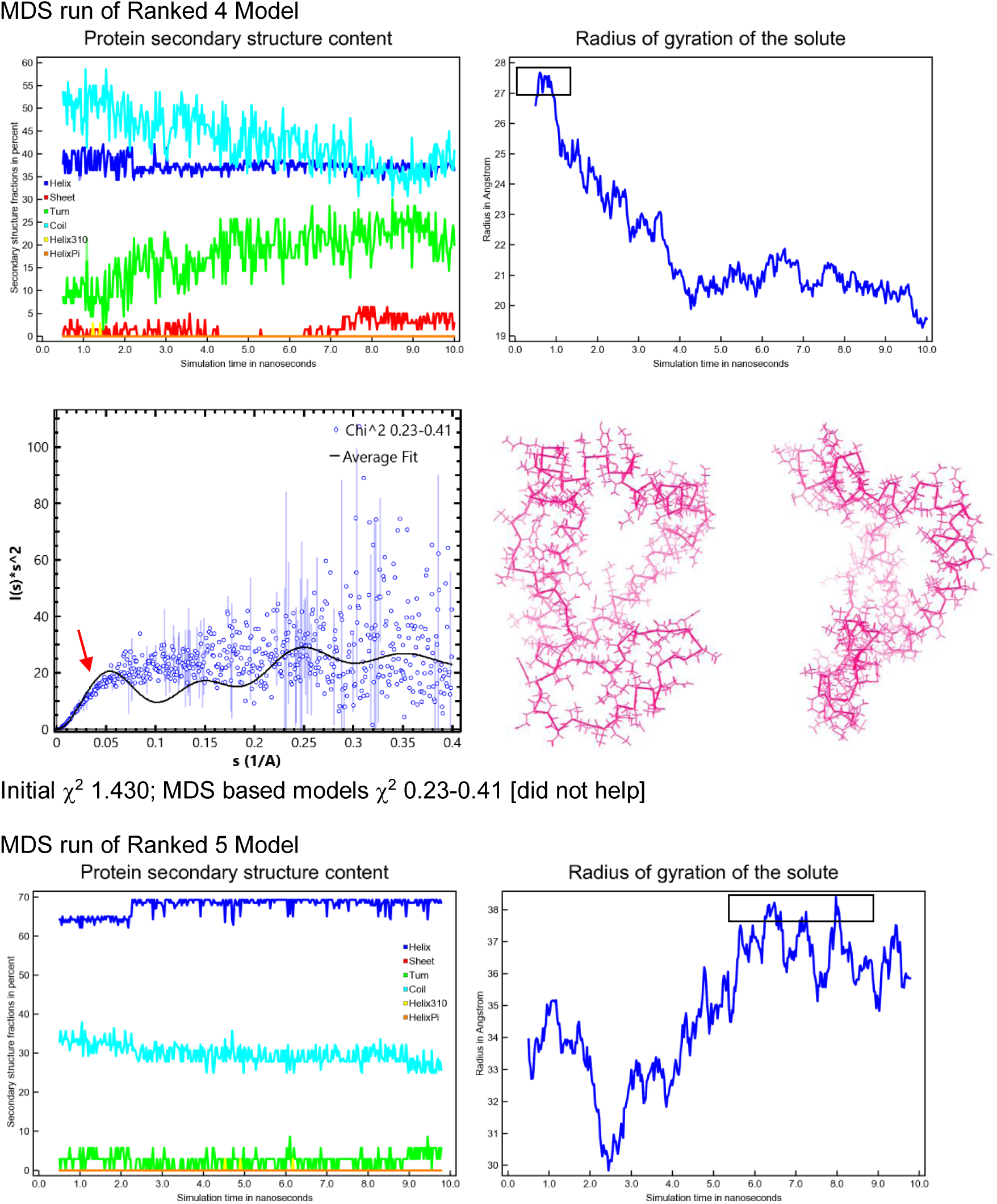
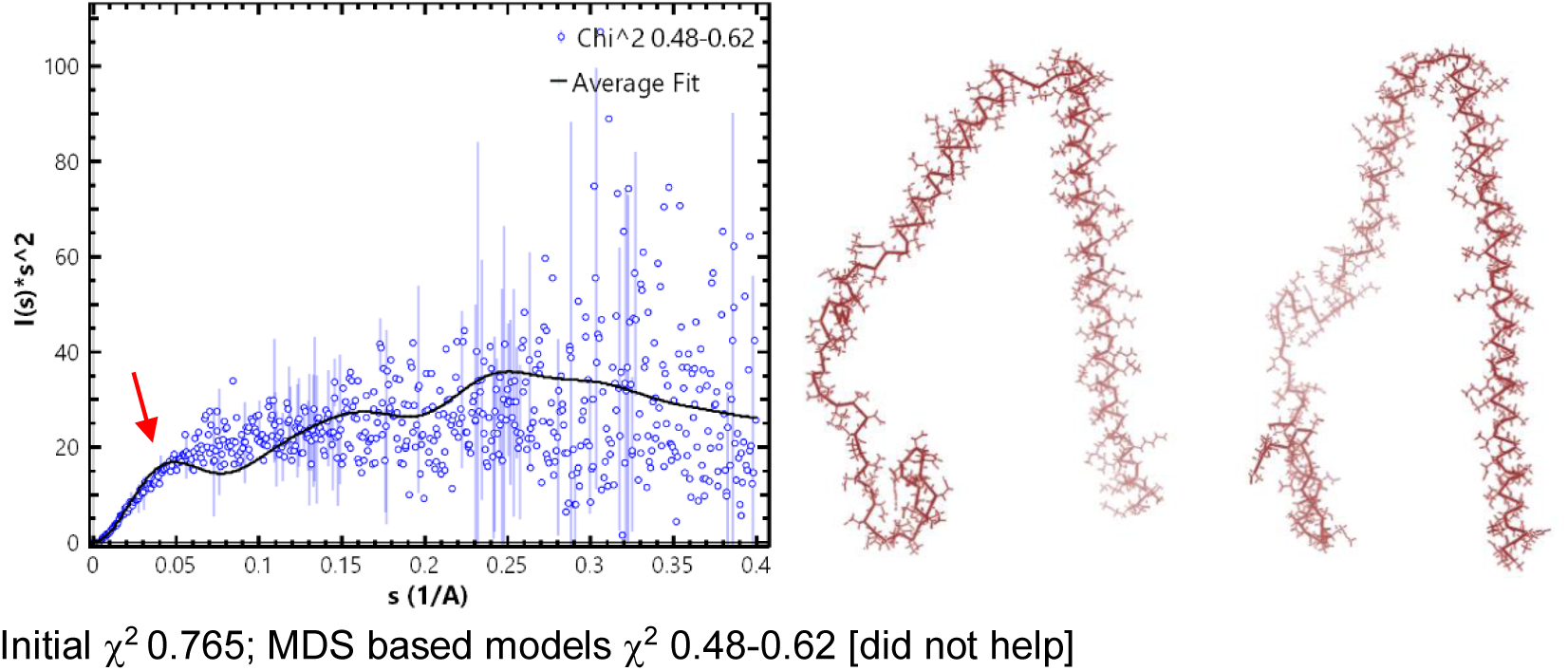
Results from MDS runs of individual predicted models are presented here. Panels show variation in secondary structural content and R_g_ values as a function of simulation time in ns. Lower set of panels show Kratky plot of experimental and theoretical SAXS profiles of selected structures from respective MDS run. Average model of selected structures are shown in two rotated views. Final χ^2^ values of the selected models vs. initial starting structure are mentioned below.

### SREFLEX Based Search

Realizing that MDS runs of individual ALPHAFOLD2 predicted models did not provide structural models which fitted both with CD and SAXS data, we applied normal mode analysis on the five models and searched for any new resultant models which better fitted the experimental SAXS data (**Fig. 8 and Supplementary Figure S7**). SREFLEX program computed different low frequency modes accessible to initial model and yielded five restrained and four unrestrained solutions for each starting structure. The computed χ^2^ values of the solutions were comparable to initial value, except for rank 4 model, where the solutions were closer to unity. For this non-extended circular shaped solution, the final χ^2^ values were 0.77-0.90 suggesting that these shapes correlate best with the acquired SAXS data. This exercise provided a larger deviation in starting structures than MDS runs which aid in providing visualization of the conformational polydispersity accessible to pre-fibrillar state of α-syn protein molecules. Normal mode analysis suggested that there is inherent motion in the terminals of the protein structure as well.

**Figure 8.**
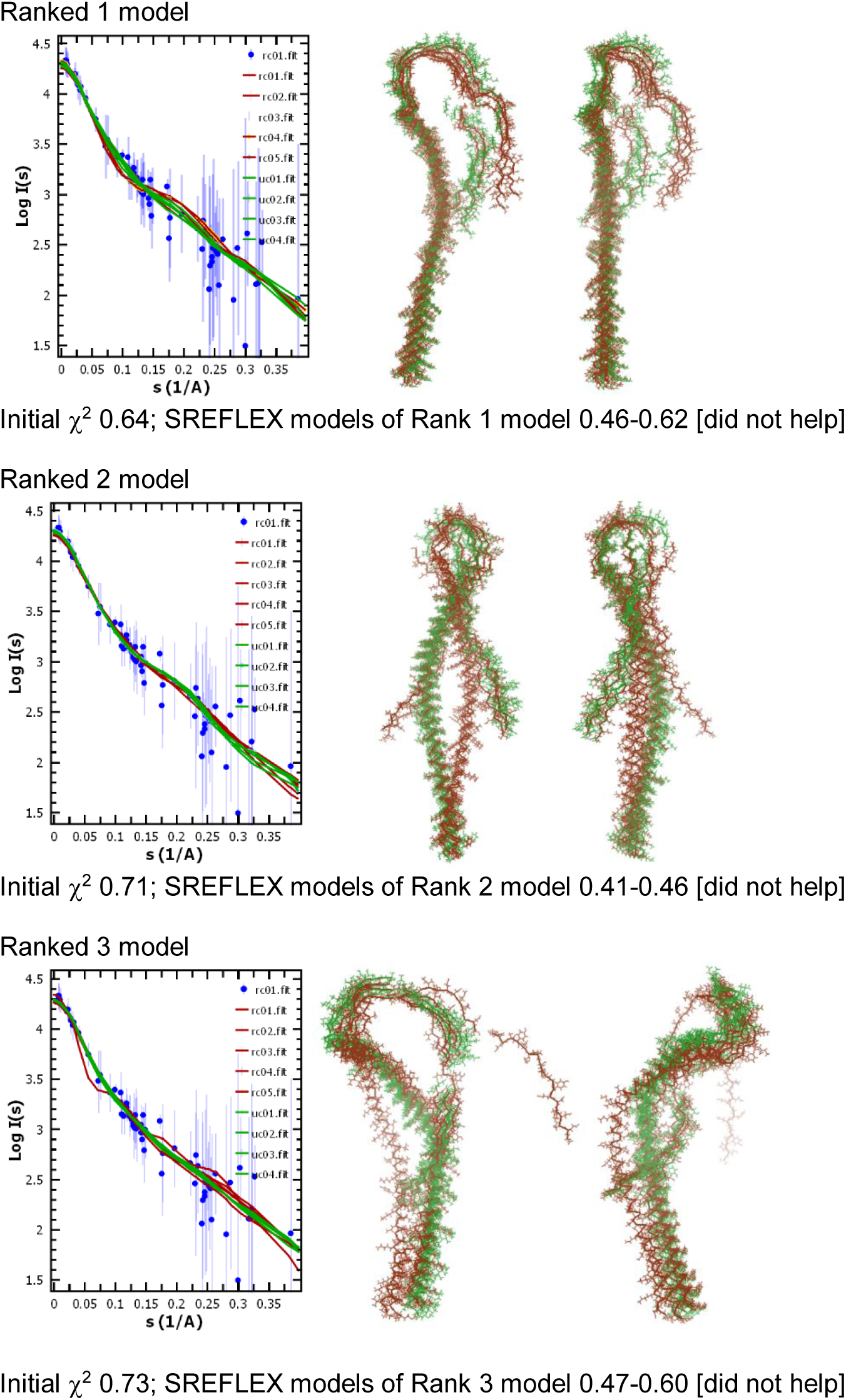
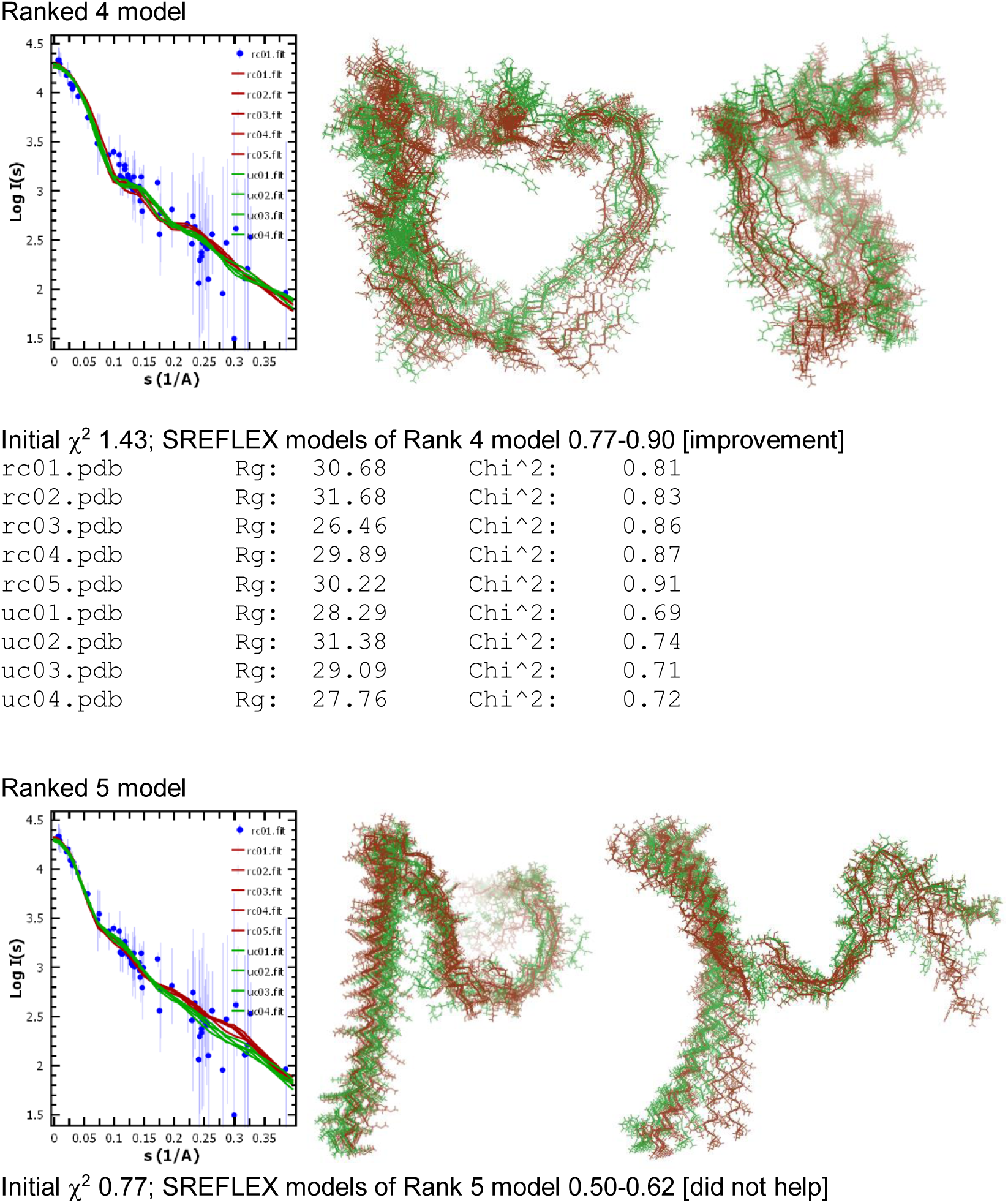
Results of attempts to search of better fitting models to SAXS data using SREFLEX program are presented here. In each set of panels, plot in left shows comparison of calculated SAXS profiles of solved models for that ranked model (lines) vs. experimental data (blue circles). Right two panels show the models solved by SREFLEX showing extent of variation possible in the ALPHAFOLD2 server based models. Unrestrained and restrained models named as uc*.pdb and rc*.pdb, and their calculated SAXS profiles, are shown in brown and forest green color, respectively. Staring χ^2^ value and range of χ^2^ values of models solved by SREFLEX are mentioned below for each model.

## Conclusion(s)

We have now carried out a detailed structural and conformational characterization of intrinsically disordered protein α-syn. CD spectroscopy suggests that recombinantly expressed and purified wild type α-syn from *E. coli.* lack any specific secondary structure as evident from spectra minimum near ∼195nm (Fauvet et al. 2012; Dong et al. 2018). The gel filtration chromatography showed that protein elutes at molecular mass higher than as expected from its monomeric state. This is consistent with previous studies showing elongated structure of α-synuclein. The ThT based fibrillation data showed that the protein propensity for fibrillation decreases significantly as temperature is decreased from 37 to 25°C which supported that the protein remains unassociated in timelines used for CD and SAXS studies at 10°C. Stretching the previous solution studies to understand the prefibrillar conformation of α-syn, we used SAXS data as reference to select the results from dummy residue modeling, EOM analysis and ALPHAFOLD2 based predictions. As reported earlier by other researchers, our SAXS data analysis also supported Gaussian-chain like scattering shape profile or signature profile of an inherently disordered protein (IDP) (**Fig. 2C**). To restore the shape of molecules in solution, in unbiased and weighted average mode, we employed Chain ensemble modeling and averaged similar models. Results were two clusters with semi-extended shapes with turns with Gaussian-Chain like scattering profiles (**Fig. 3B and C**). Extending our analysis further, we employed EOM calculations which used primary structure of protein during search of templates and SAXS data was used as search reference. Two runs were done, one considering disorder across whole length of protein and second biasing first 60 residues as α-helix. Results shown in **Figs. 4B and 5B** show mole fractions of different conformations which should compose back the experimental scattering data. Interestingly, one model solved using EOM i.e. 7083 showed χ^2^ value of 1 which meant best resemblance to experimental data albeit it had lower mole fraction. Either case showed that upon aligning the N-terminal of different conformations, the central and C-terminal part of molecule wiggled substantially in space giving view on the disordered state of the protein.

To bring in the new aspect of ALPHAFOLD2 server-based predications, we used sequence of wild type α-syn molecule to assess predicted structures vs. SAXS data. To the best of our knowledge, the images provided in **Fig. 6** are the first all-atom models of α-syn predicted from ALPHAFOLD2 server and then compared with experimental data from solution. We obtained models with extended shapes with a U-turn, and curved models, all with comparable deviation from unity in their computed χ^2^ values. Their overlap also agreed to conclusions made from EOM analysis that if N-terminal is superimposed then the C-terminal wobbles in space, but could not provide cue on mole fraction existence in solution. Additionally, the R_g_ values of models predicted by ALPHAFOLD2 server were substantially smaller than experimental value of 47 Å and EOM analysis. To explore better convergence of the χ^2^ value to SAXS data, we performed MDS runs and normal mode analysis on ALPHAFOLD2 results. Only for the curved model (rank 4) preferred secondary structural content similar to CD data analysis but its R_g_ values remained consistently below experimentally determined value. Selected structures from MDS runs for top 3 ranked models were comparable to experimental R_g_ value, but had significant molecular structure as α-helix which had opened up in space than starting hairpin like predicted structure during simulation. Based on R_g_ and χ^2^ values, models froom MDS run for 3^rd^ ranked model seemed better fitting to SAXS data. Selected models from MDS runs for 2^nd^ and 3^rd^ ranked models resembled EOM run based model 7083 which best agreed with experimental data.

To push more structural changes than possible by MDS runs, we performed normal mode analysis to search for structures with χ^2^ values closer to unity. Only ranked 4 model with curved shape led to models with final best resemblance to experimental data. Interestingly, this model’s MDS runs also matched with CD data. Final results of SREFLEX program on this model suggests that if this model opens up in space (instead of collapsing as in MDS), it can provide good representation of ensemble in CD and SAXS data. Our overall analysis concludes that a range of extended, bent, distorted and curved structures are simultaneously accessible to α-syn protein in solution. Re-searching of conformational space using SAXS data as reference suggested that a curved model may dominate prefibrillar structure of wild type α-syn. This observation is in agreement to previous NMR data based indirect interpretations. Since, fibrillation rate is dependent on multitude of factors like protein sequence, concentration of protein, ionic strength/pH of buffer, presence of osmolytes, storage time/temperature and some other factors, deciphering prefibrillar conformation and its commitment to native fibril formation will require different repetitions of experiments like ours. We paved a way to understand molecular shapes accessible to α-syn under a set of conditions which is definitely more reliable than unbiased/steered MD simulations, and easier than NMR data-based restorations. Due to the labile nature of this small protein, CryoEM and/or crystallography can be ruled out to provide true insight into α-syn in monomeric state. Global insight into conformations accessible to α-syn is highly sought to embark on computer-aided drug discovery to retard the process of fibrillation of this protein and other IDPs. While, we will continue to re-apply the protocol mentioned here on other variants of wild type α-syn, we invite other researchers to follow our suite.

## Acknowledgements

Authors acknowledge funding support from UNSEEN project from 12FYP of CSIR INDIA which established the SAXS facility in CSIR-IMTECH. This is IMTECH communication number 50/2023.

