## Supplementary figures for "VISUALIZING GAUSSIAN-CHAIN LIKE STRUCTURAL MODELS OF HUMAN α-SYNUCLEIN IN MONOMERIC PRE-FIBRILLAR STATE: SOLUTION SAXS DATA AND MODELING ANALYSIS"


**Supplementary Figure 1** Used primary α-syn primary structural details are presented here. (A) KEGG link , (B) uniport ID for human α-syn sequence used in this study are mentioned here. (C) Schematic representation of primary α-syn structure represents the amino acid sequence used in this study.

**Supplementary Figure 2** Additional results for α-syn purification and thioflavin-T mediated fluorescence kinetics are presented here. (A) Protein purification profile using Gel Filtration Chromatography is shown here. (B) Elution profiles of standard proteins used for $K_{av}$ calculation are plotted here. (C) Image of 15% SDS-PAGE purification profile of α-syn done is shown here. Left most lane is of loaded sample (lane 1), next to right (lane 2) is of the collected fraction from peak at 16 mL. Lane M represents the used molecular weight marker (BioRad #1610374) for comparison. (D) *In-vitro* α-syn fibrillation at 25°C with lysozyme as non-fibrillatory negative control is presented**.**

**Supplementary Figure 3** Additional results from EOM analysis considering all 140 residues to be disordered are presented here. (A) Statistics of the computed flexibility of the ensemble pool used for search are mentioned. (B) Upper left plot shows the fit of the computed SAXS profile of the solved models (red line) vs. experimental data (blue dots). Lower left panel shows the fit residuals of above comparison. Upper and lower right panels show the distribution of the $D_{max}$ and $R_g$ values of the models in the selected pool (grey histogram), and the blue histogram indicate frequency and models which were considered to be compared with experimental SAXS data and sequence given, respectively.

**Supplementary Figure 4** Additional results from EOM analysis considering first 60 residues to be adopting helical order and rest of the residues adopting disordered structure. Rest descriptions are analogous to those described for **Fig. S3**.

**Supplementary Figure 5** Additional results from ALPHAFOLD2 server based predictions are shown here. (A) Predicted local Distance Difference Test (lDDT) values per residue of $\alpha$Syn sequence are plotted here for the five top ranked models. (B) Predicted Aligned Error (PAE) values for the five ranked 3D models of $\alpha$Syn protein are shown here. (C) Number of sequences covered with their identity vs. the query sequence are plotted here.

**Supplementary Figure 5** Additional results from ALPHAFOLD2 server based predictions are shown here.

**Supplementary Figure 6** Per residue secondary structural content as a function of simulation and overall RMSF values are shown here for the top 5 ranked models predicted by ALPHAFOLD2 server.

**Supplementary Figure 7** Details of SREFLEX program based searches for better fitting models are shown here with top 5 ranked models from ALPHAFOLD2 server as initial structures.

**Supplementary Figures**

**Supplementary Figure S1**

A. The link to find the α-syn sequence, using KEGG:  https://www.genome.jp/entry/hsa:6622

B. UniProt ID:  P37840

C. Primary structure of α-syn

**MDVFMKGLSKAKEGVVAAAEKTKQGVAEAAGKTKEGVLYVGSKTKEGVVHGVATVAEKTEQVTNVG GAVVTGVTAVAQKTVEGAGSIAAATGFVKKDQLGKNEEGAPQEGILEDMPVDPDNEAYEMPSEEGYQ DYEPEA**

N' 1-60 residues are highlighted in blue, NAC region 61-95 residues are highlighted in orange and C' residues are highlighted in green.

**Supplementary Figure S2**

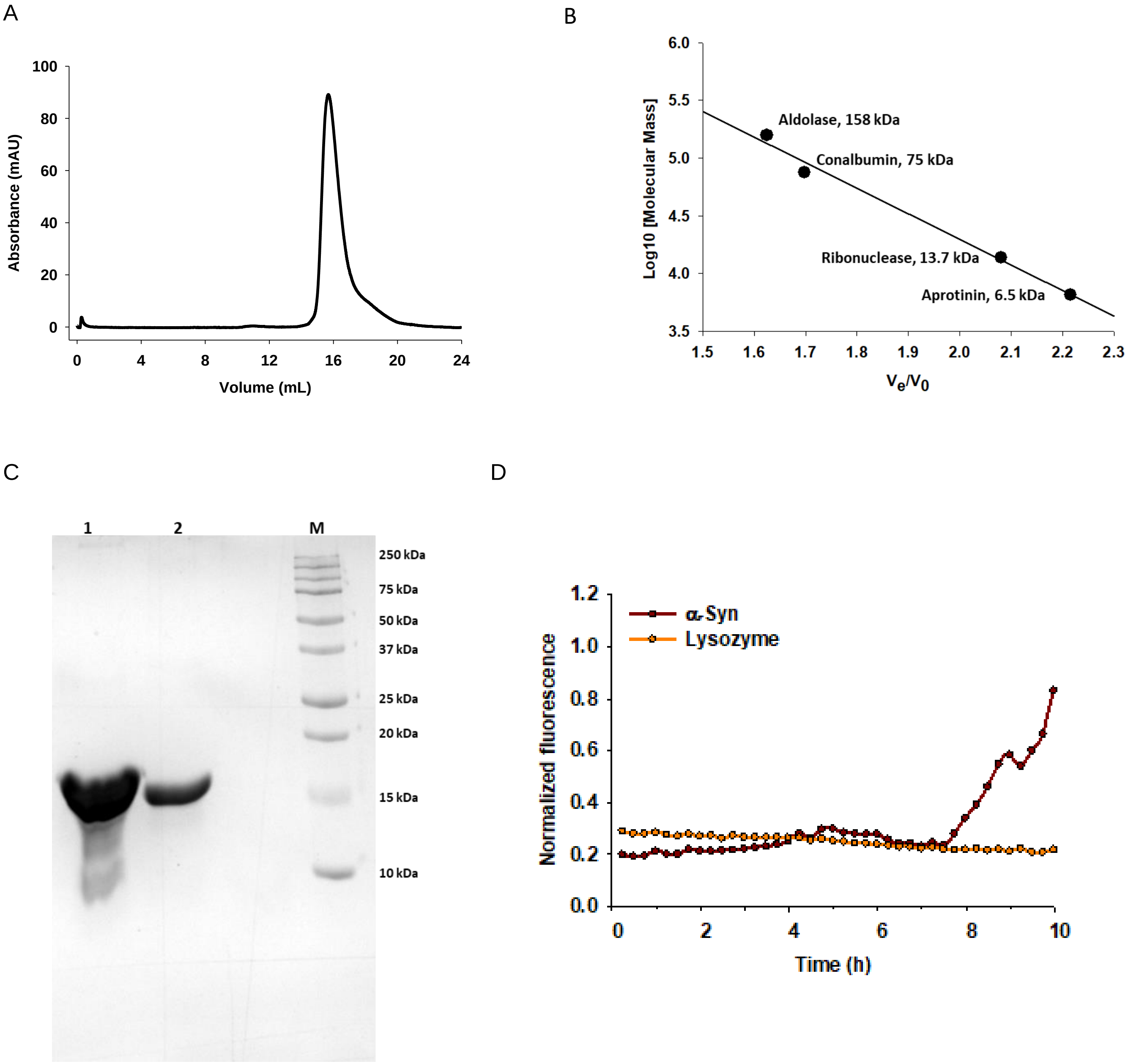

**Supplementary Figure S3**

EOM Results: Considering all 140 residues to be disordered

A Statistics of computed flexibility

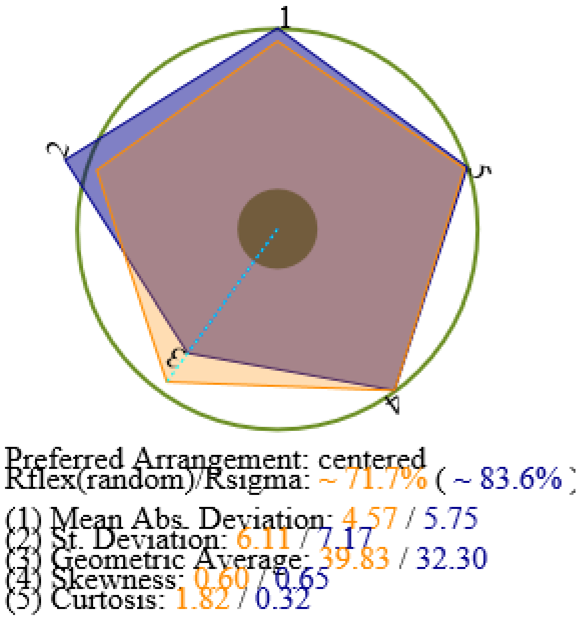


B

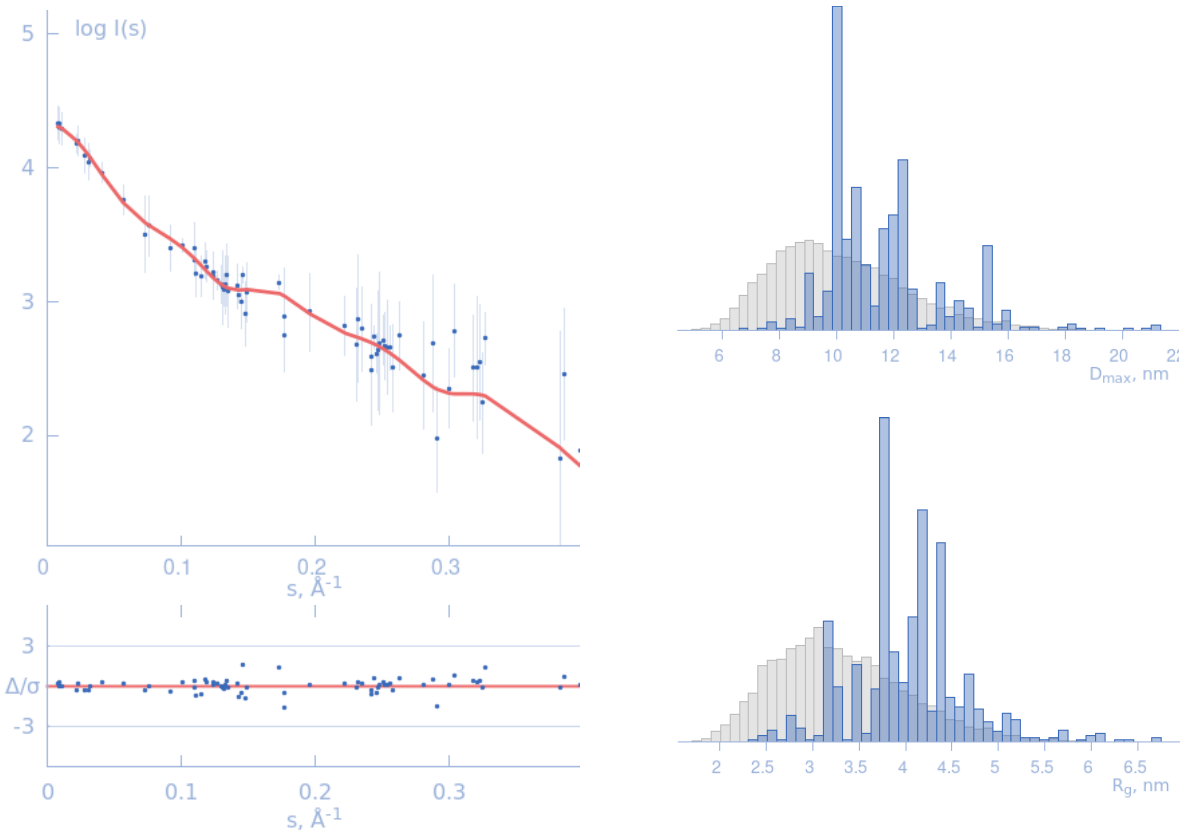

**Supplementary Figure S4**

EOM Results: Considering first 60 residues to be helical and remaining residues to be disordered

A  Statistics of computed flexibility

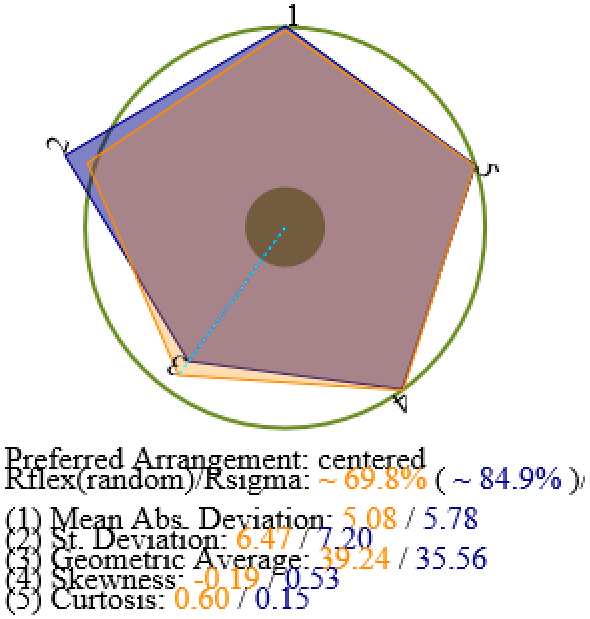


B

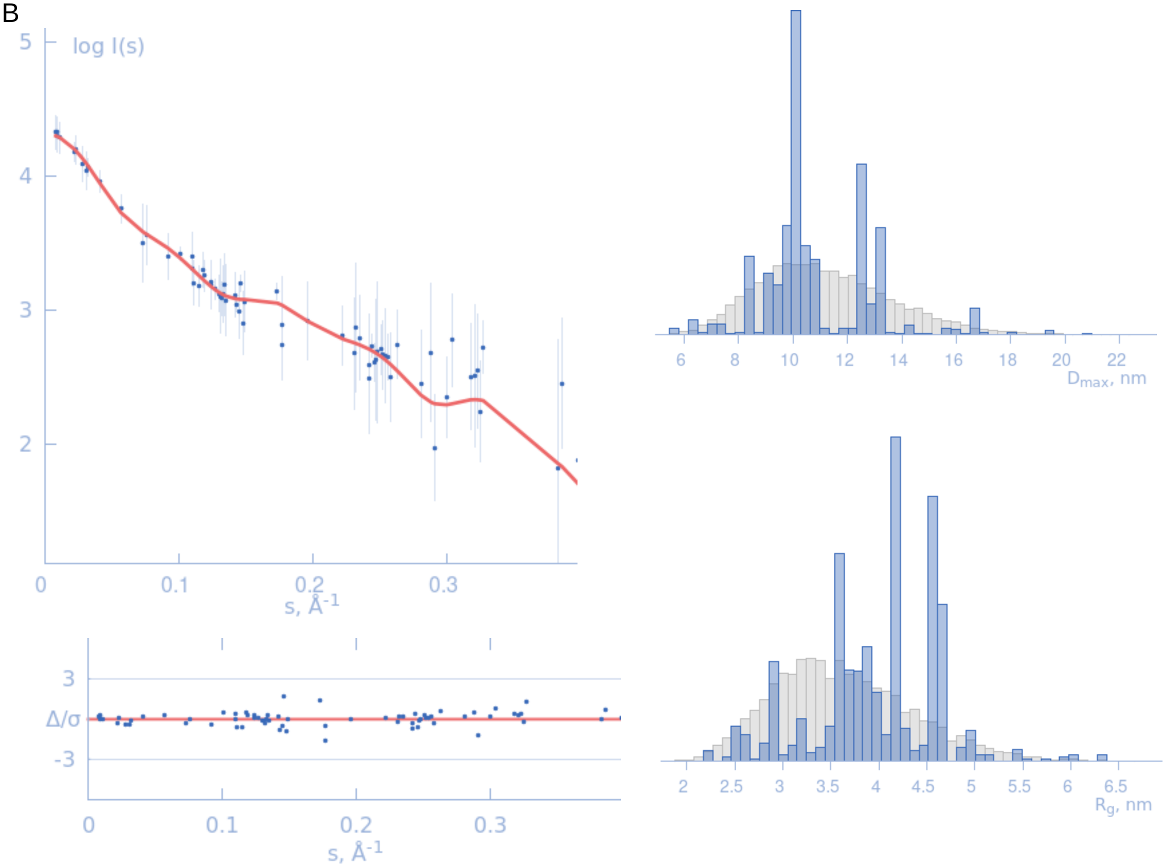

**Supplementary Figure S5**

A

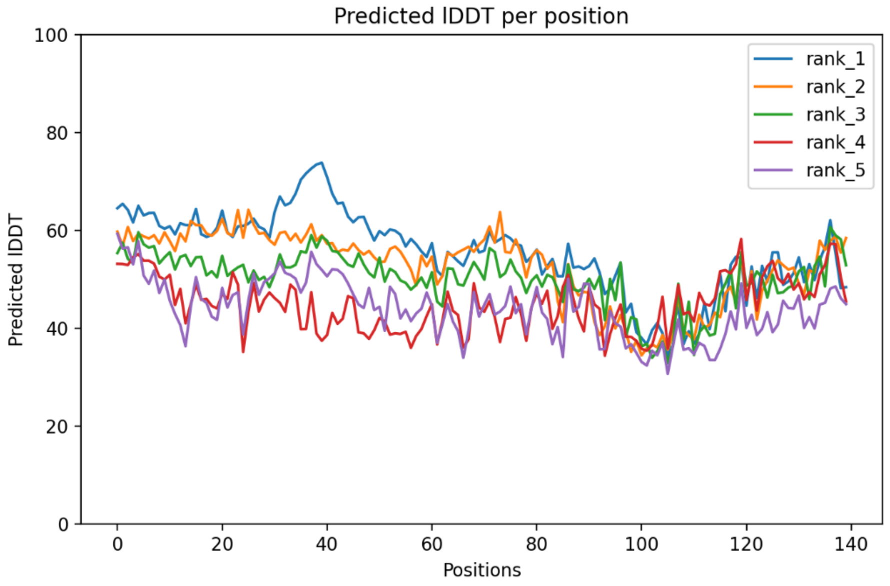


B

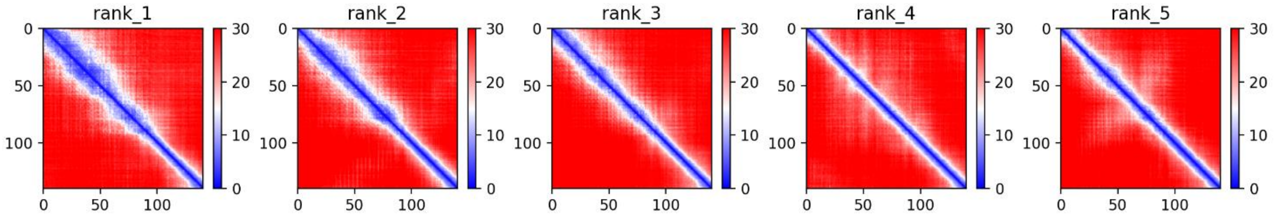


C

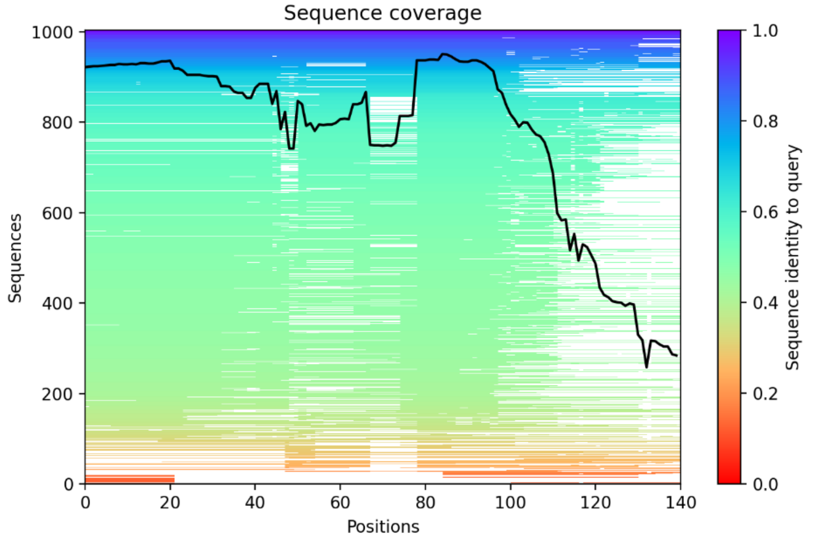

**Supplementary Figure S6**

### Ranked 1 Model

Per-residue protein secondary structure of molecule A

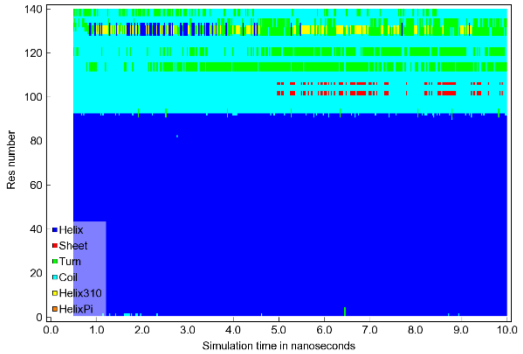


Solute protein/nucleic acid residue RMSF

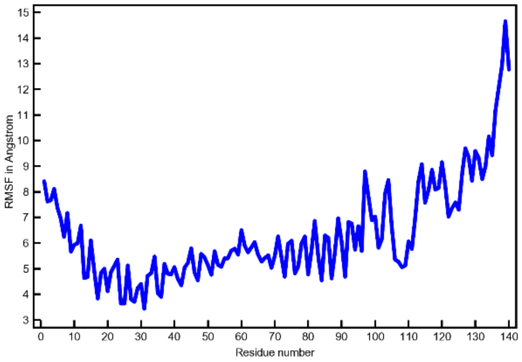


### Ranked 2 Model

Per-residue protein secondary structure of molecule A

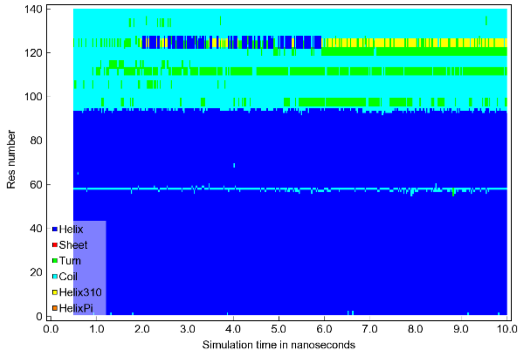


Solute protein/nucleic acid residue RMSF

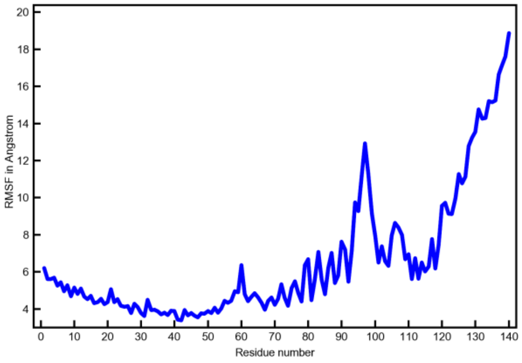


### Ranked 3 Model

Per-residue protein secondary structure of molecule A

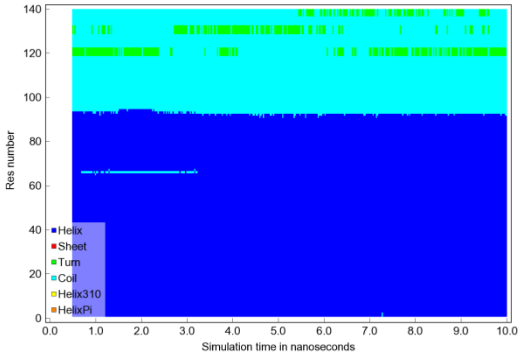


Solute protein/nucleic acid residue RMSF

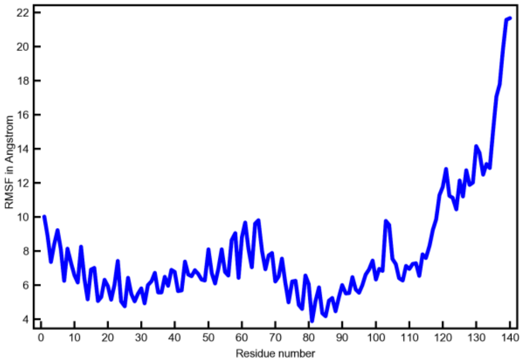

### Ranked 4 Model

Per-residue protein secondary structure of molecule A

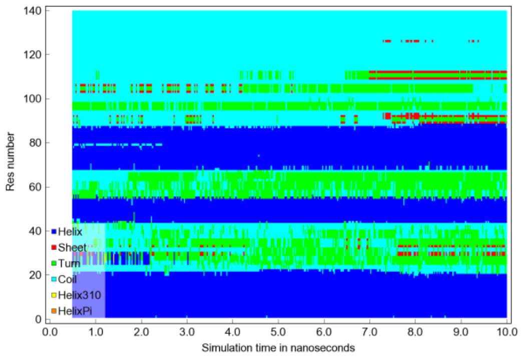


Solute protein/nucleic acid residue RMSF

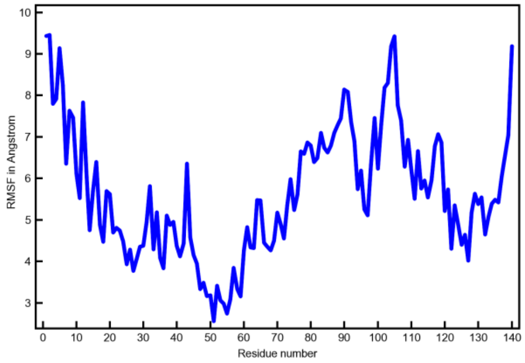


### Ranked 5 Model

Per-residue protein secondary structure of molecule A

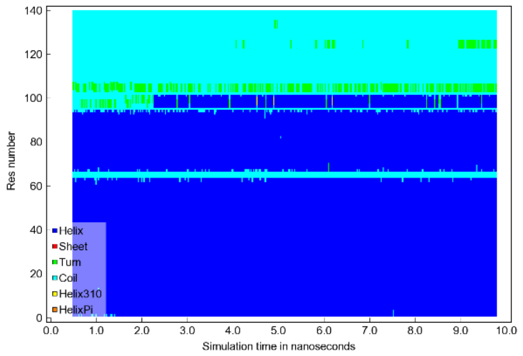


Solute protein/nucleic acid residue RMSF

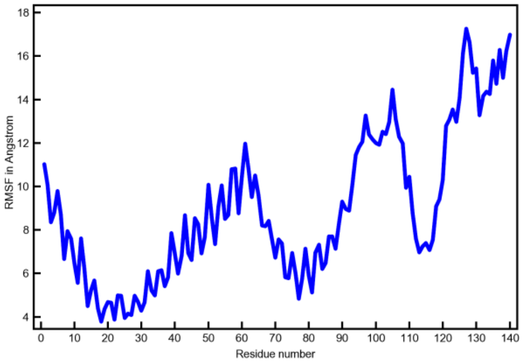

**Supplementary Figure S7**

SREFLEX Results on AlphaFold2 Solutions

```
Rank1
--SREFLEX REPORT
--VERSION: ATSAS 3.0.0 (r11734)
--REFERENCE: Panjkovich & Svergun (2016) PCCP 18, 5707-19
--DATE AND TIME of run: 2022-11-02 19:30:59
--INPUT SAS data filename: syn_30x3_8mg_003Frames_corrected[5].dat
--INPUT PDB coordinates: r1m1.pdb
--Initial Chi2: 0.64
--Initial clashes: 0
--RESULTS SUMMARY:
model   Chi2   RMSD  breaks  clashes  modes
rc01    0.51  11.53  0.33    0.00     n7s-5xn10s1
rc02    0.61   2.93  0.01    0.00     n10s1
rc03    0.61   9.51  0.13    0.00     n7s-4
rc04    0.61   2.53  0.00    0.00     n7s-1
rc05    0.62   2.29  0.03    0.00     n13s1
uc01    0.46  11.23  0.81    0.00     n7s-5xn10s1_X_n10s2xn16s-1
uc02    0.47   9.51  0.41    0.00     n7s-4_X_n10s-2xn11s-2
uc03    0.49   9.72  0.84    0.00     n7s-4_X_n8s2xn12s1
uc04    0.50   4.78  0.01    0.00     n7s-1_X_n9s2xn10s2
--RUNTIME: 2.07 minutes, 8 CPU cores/threads.

Rank2
--SREFLEX REPORT
--VERSION: ATSAS 3.0.0 (r11734)
--REFERENCE: Panjkovich & Svergun (2016) PCCP 18, 5707-19
--DATE AND TIME of run: 2022-11-02 19:33:03
--INPUT SAS data filename: syn_30x3_8mg_003Frames_corrected[5].dat
--INPUT PDB coordinates: r2m4.pdb
--Initial Chi2: 0.71
--Initial clashes: 0
--RESULTS SUMMARY:
model   Chi2   RMSD  breaks  clashes  modes
rc01    0.45  12.15  0.91    0.00     n8s-4xn12s-5
rc02    0.45  11.40  0.02    0.14     n8s-4xn13s2
rc03    0.46  10.18  0.64    0.00     n8s-4xn14s-5
rc04    0.46  13.94  0.69    0.00     n7s-5xn9s2
rc05    0.46  12.00  0.45    0.00     n7s-4xn8s3
uc01    0.40  11.40  0.74    0.00     n8s-4xn14s-5_X_n7s-1xn9s-1
uc02    0.41  12.47  0.41    0.14     n8s-4xn13s2_X_n13s-1xn16s-2
uc03    0.41  10.39  0.42    0.00     n8s-4xn14s-3_X_n7s1xn12s2
uc04    0.41  10.19  0.76    0.00     n8s-4xn14s-5_X_n7s-1xn10s-2
--RUNTIME: 4.82 minutes, 8 CPU cores/threads.

Rank3
--SREFLEX REPORT
--VERSION: ATSAS 3.0.0 (r11734)
--REFERENCE: Panjkovich & Svergun (2016) PCCP 18, 5707-19
--DATE AND TIME of run: 2022-11-02 19:37:53
--INPUT SAS data filename: syn_30x3_8mg_003Frames_corrected[5].dat
--INPUT PDB coordinates: r3m2.pdb
--Initial Chi2: 0.73
--Initial clashes: 0
```

```
--RESULTS SUMMARY:
model   Chi2   RMSD  breaks  clashes  modes
rc01    0.48  12.47  0.29    0.00     n7s-3xn10s3
rc02    0.51  15.04  0.87    0.14     n7s-5xn16s1
rc03    0.53  14.89  0.89    0.29     n7s-5xn12s1
rc04    0.57  14.76  0.51    0.14     n7s-5
rc05    0.60   8.93  0.07    0.00     n7s3
uc01    0.47  15.18  0.82    0.14     n7s-5_X_n11s1xn15s2
uc02    0.48  15.26  0.83    0.29     n7s-5xn12s1_X_n11s-1xn14s1
uc03    0.48  12.45  0.86    0.43     n7s-4_X_n13s1xn14s-2
uc04    0.48  15.62  0.95    0.29     n7s-5xn12s1_X_n11s-2xn14s1
--RUNTIME: 3.32 minutes, 8 CPU cores/threads.

Rank4
--SREFLEX REPORT
--VERSION: ATSAS 3.0.0 (r11734)
--REFERENCE: Panjkovich & Svergun (2016) PCCP 18, 5707-19
--DATE AND TIME of run: 2022-11-02 19:41:14
--INPUT SAS data filename: syn_30x3_8mg_003Frames_corrected[5].dat
--INPUT PDB coordinates: r4m3.pdb
--Initial Chi2: 1.43
--Initial clashes: 0
--RESULTS SUMMARY:
model   Chi2   RMSD  breaks  clashes  modes
rc01    0.81   8.09  0.09    0.00     n9s-2xn10s2
rc02    0.86  10.39  0.22    0.00     n9s-3xn10s2
rc03    0.87   8.98  0.89    0.43     n7s3xn11s-2
rc04    0.88   6.22  0.07    0.00     n9s-1xn10s2
rc05    0.90   6.56  0.01    0.00     n9s-2xn10s1
uc01    0.76   8.83  0.62    0.43     n7s3xn11s-1_X_n10s2xn11s-2
uc02    0.77   8.63  0.09    0.00     n9s-2xn10s1_X_n8s-2xn14s2
uc03    0.77   8.15  0.49    0.00     n7s2xn11s-2_X_n8s-2xn10s-2
uc04    0.77   8.64  0.54    0.43     n7s3xn11s-1_X_n10s1xn11s-2
--RUNTIME: 6.33 minutes, 8 CPU cores/threads.

Rank5
--SREFLEX REPORT
--VERSION: ATSAS 3.0.0 (r11734)
--REFERENCE: Panjkovich & Svergun (2016) PCCP 18, 5707-19
--DATE AND TIME of run: 2022-11-02 19:47:36
--INPUT SAS data filename: syn_30x3_8mg_003Frames_corrected[5].dat
--INPUT PDB coordinates: r5m5.pdb
--Initial Chi2: 0.77
--Initial clashes: 0
--RESULTS SUMMARY:
model   Chi2   RMSD  breaks  clashes  modes
rc01    0.59   8.10  0.51    0.00     n8s2xn10s2
rc02    0.60   6.48  0.26    0.00     n8s2xn10s1
rc03    0.61  14.63  0.73    0.00     n7s-5xn10s2
rc04    0.62   6.11  0.35    0.00     n8s1xn10s2
rc05    0.62   9.78  0.37    0.00     n7s-3xn10s2
uc01    0.50   8.66  0.74    0.00     n7s-2xn10s2_X_n15s-2xn16s2
uc02    0.51   8.85  0.74    0.00     n8s2xn10s2_X_n14s2xn16s2
uc03    0.51   8.80  0.61    0.00     n8s2xn10s2_X_n16s2
uc04    0.52   7.24  0.30    0.00     n8s2xn10s1_X_n16s2
--RUNTIME: 5.30 minutes, 8 CPU cores/threads.
```
